# Zebrafish arterial valve development occurs through direct differentiation of second heart field progenitors

**DOI:** 10.1101/2024.04.17.589935

**Authors:** Christopher J. Derrick, Lorraine Eley, Ahlam Alqahtani, Deborah J. Henderson, Bill Chaudhry

## Abstract

**Aims:** Bicuspid Aortic Valve (BAV) is the most common congenital heart defect, affecting at least 2% of the population. The embryonic origins of BAV remain poorly understood, limiting the identification of assays for validating patient variants and ultimately causative genes for BAV. In both human and mouse, the left and right leaflets of the arterial valves arise from the outflow tract cushions, with interstitial cells originating from neural crest cells and endocardial-to-mesenchymal transition (EndoMT). In contrast, an EndoMT-independent mechanism of direct differentiation by cardiac progenitors from the second heart field (SHF) is responsible for the formation of the anterior and posterior leaflets. Defects in either of these developmental mechanisms can result in BAV. Although zebrafish have been suggested as a model for human variant testing, their naturally bicuspid arterial valve has not been considered suitable for understanding human arterial valve development. Here, we have set out to investigate to what extent the processes involved in arterial valve development are conserved in zebrafish and ultimately, whether functional testing of BAV variants could be carried out in zebrafish.

**Methods and Results:** Using a combination of live imaging, immunohistochemistry and Cre-mediated lineage tracing, we show that the zebrafish arterial valve primordia develop directly from undifferentiated SHF progenitors with no contribution from EndoMT or neural crest, in keeping with the human and mouse anterior and posterior leaflets. Moreover, once formed, these primordia share common subsequent developmental events with all three mammalian arterial valve leaflets.

**Conclusions:** Our work highlights a conserved ancestral mechanism of arterial leaflet formation from the SHF and identifies that development of the zebrafish arterial valve is distinct from that of the atrioventricular valve. Crucially, this confirms the utility of zebrafish for understanding the development of specific BAV subtypes and arterial valve dysplasia, offering potential for high-throughput variant testing.

**Translational Perspective:** Large genomic studies of patients with Bicuspid Aortic Valve (BAV) have identified numerous variants predicted to be causative, yet due to a lack of suitable, *in vivo* functional assays, advancement of genetic testing, discussion of risk to family members and accurate prognosis is not yet widely possible. Here, we show that zebrafish demonstrate a high level of conservation in arterial valve development with the intercalated leaflets in human, establishing zebrafish as a suitable *in vivo* model that can begin to overcome the disconnect between clinical genetics and developmental biology.

## Introduction

The aortic and pulmonary valves ensure unidirectional blood flow from the left and right ventricles to the systemic and pulmonary circulations respectively. These valves can be impacted by congenital malformations, most frequently Bicuspid Aortic Valve (BAV), which occurs in at least 2% of the population, typified by the presence of two valve leaflets rather than three^1,2^. BAV can be present in isolation, but is also associated with other congenital heart malformations, such as Hypoplastic Left Heart Syndrome^3^ or chromosomal disorders such as Down’s Syndrome^4^. Although frequently undetected at birth, BAV predisposes individuals to a spectrum of progressive aortic abnormalities as well as degeneration of the valve with age, leading to stenosis or regurgitation and ultimately ventricular failure if untreated^5^. Defects of the pulmonary valve are less common but are strongly linked with other heart malformations such as Tetralogy of Fallot^3,6^.

The semi-lunar valves develop in the arterial roots, located at the boundary between the ventricular myocardium and the smooth muscle of their respective arterial trunks, which in amniotes has a characteristic structure: hinge points attaching the leaflets to the myocardial wall, sinuses and fibrous interleaflet triangles^7,8^. Leaflet structure broadly conserved across vertebrates, with apparently distinct elastin and collagen rich layers^9,10^. Despite the importance of the arterial valves, remarkably little is known about the developmental mechanisms that give rise to their leaflets and until recently, studies of the atrioventricular valves had been extrapolated to understand arterial valve development ^11^.

Addition of multipotent cardiac progenitors from the second heart field (SHF) is critical for normal arterial pole development^12,13^. These cells express the transcription factor Islet1 (Isl1), which is downregulated upon addition to the heart At the arterial pole, the first wave of SHF cells form cardiomyocytes^14,15^ and later smooth muscle^7,16^. Within the distal outflow tract, cells pass through the transition zone where they co-express Isl1 and mature cardiomyocyte markers^14,15^; this region eventually forms the arterial root. At embryonic day (E) 10.5-E11.5 in mice (Carnegie stage (CS) 14-16 in humans), two tongues of Isl1+ cells, that neither express markers of myocardial nor smooth muscle identity, can be identified in the wall of the still unseptated outflow tract^14,17,18^. The most proximal of these undifferentiated cells form the intercalated valve swellings (ICVS)^14,17,18^ and the remainder differentiate later into the smooth muscle of the arterial walls. Concomitant with ICVS formation, the two cardiac jelly-rich outflow tract (OFT) cushions, which are found along the entirety of the (proximal) myocardial part of the outflow tract, are populated by cells from the overlying endocardium (itself SHF-derived) through Endothelial-to-Mesenchymal transition (EndoMT) and from the cardiac neural crest^17–19^. These cushions expand and ultimately fuse, leading to outflow tract septation^20,21^. The distal ends of these main cushions form the right and left leaflets of the aortic and pulmonary valve, whereas the ICVSs form the anterior and posterior/non-coronary leaflets^17^. The different origins of these leaflets is reflected in the cell lineages they contain, as shown by Cre-based lineage tracing performed in mouse. The left and right leaflets – derived from the outflow cushions – are composed of valve interstitial cells (VICs) originating from EndoMT and neural crest cells lineages. In contrast, the anterior and non-coronary leaflets – derived from the ICVS – have few EndoMT- and neural crest cell-derived VICs and instead are formed from a distinct lineage of SHF progenitors, without passing through the endocardial lineage^17^. All mesenchymal cells within these six primordia express the transcription factor Sox9^18,22^. Notably however, the cells in the ICVS differentiate directly from Isl1+ SHF progenitors into Sox9-expressing VICs, where these cells co-express both markers for a short period^17^, not seen in the OFT cushion-derived leaflets. A similar pattern of co-expression of ISL1 and SOX9 is visible in the developing ICVS of human embryos between CS13-16, supporting this is a conserved mechanism of primordia formation^18^. Despite these distinct origins, once established, all primordia undergo the same remodelling processes. The understanding of these distinct mechanisms of primordia formation may reflect the relationship between BAV subtypes^1,23^ and aortopathies. Therefore, there is need for an understanding of the relationship between OFT development, patient mutations and their associated disease to identify the developmental origins of arterial pole malformations such as BAV.

Animal models have been invaluable in understanding the complex processes of valve formation, traditionally mouse, and more recently zebrafish. The rapid, temporally reproducible development of zebrafish is highly amenable to *in vivo* live imaging and, together with a high level of conservation of both process and gene regulatory networks, has made them a powerful model to investigate aspects of cardiac development not possible in mouse^24^. The zebrafish has a single, naturally bicuspid arterial valve, residing in the unseptated outflow tract between the single ventricle and the bulbous arteriosus (BA)^25,26^. This has led many to assume that it cannot be used to model human arterial valve development or disease. However, the BA is a smooth muscle, elastin-rich structure^26^, and both the BA and distal part of the ventricle are derived from the SHF^13,27^, resembling the amniote arterial root. In the few studies that have characterised the arterial valve in zebrafish^28–31^, the origin of the leaflets and whether they have any similarities to human arterial valve development have not been carried out and, similar to mouse, many others have made assumptions as to the similarity of their development with the atrioventricular valve leaflets^32,33^. Establishing the mechanism by which the arterial valve forms, if similar to mouse/human, would enable zebrafish to be a useful model for investigating key aspects of arterial pole development and disease.

In this study, we ask to what extent the processes involved in arterial valve development are conserved in zebrafish. We investigated whether the zebrafish arterial root and leaflets are structurally similar to those in the mouse/human; whether the zebrafish possesses a transition zone, and if this the site of arterial valve formation. We then examined whether the zebrafish arterial valve primordia develop through cellularisation of a cushion or through direct differentiation of SHF progenitors. By identifying the conserved developmental mechanisms leading to the formation of zebrafish arterial valve primordia, we establish the suitability of zebrafish for understanding the development of specific BAV subtypes and arterial valve dysplasia, and as a tool for patient variant analysis.

## Methods

### Zebrafish husbandry

Adult zebrafish (*D. rerio*) were maintained according to standard laboratory conditions and all procedures carried out in accordance with the local Animal Welfare and Ethical Review Body (AWERB), UK Home Office and Newcastle University (Project Licences P25F4F0F4 and PP0696166) in line with Directive 2010/63/EU of the European Parliament. Adult zebrafish were euthanised by terminal anaesthesia using Tricaine methanesulfonate (MS222, Merck E10521) followed by destruction of the brain. Details of specific lines used, collection of embryonic and adult tissue are listed in supplemental methods.

### Mouse tissue

WT C57BL/6 mice were maintained according to standard laboratory conditions and all procedures carried out in accordance with the local Animal Welfare and Ethical Review Body (AWERB) and Newcastle University (Project Licences 30/3876 and P9E095FF4) in line with Directive 2010/63/EU of the European Parliament. At P90, the animal was culled by cervical dislocation, the heart excised, the region of interest dissected, fixed in 4% PFA, serial dehydrated and embedded in paraffin wax (VWR, 361077E) following standard protocols. Wax sections were cut at 8μm thick using a Leica RM2235 Microtome, with alternating sections split between three different groups for analysis of elastin, proteoglycans and collagen within the same heart.

### Live imaging

*Tg(kdrl:GFP)* embryos were imaged in 2% Methyl Cellulose (Sigma M0262) on an Olympus BX61 microscope using cellSens Dimension software (Evident).

### BrdU incorporation

BrdU (Merck, B5002) was dissolved in E3 at a concentration of 10mg/mL, filtered using a 0.22um filter, aliquoted and stored at -20C for long term storage. For BrdU pulses, 20 manually dechorionated *Tg(kdrl:GFP)* embryos were incubated in 3mL of 5mg/mL BrdU in E3 at 28.5C. Embryos were rinsed in E3 and fixed in 4% PFA overnight at 4C then dehydrated to 100% MeOH and stored at -20C.

Following rehydration and prior to immunohistochemistry, embryos were incubated in 2N HCl for 1 hour at room temperature, and were then extensively washed prior to blocking (see below)

### Paraffin embedding of adult zebrafish tissue

For Miller’s Elastin, Alcian Blue and Masson’s Trichrome, adult hearts were washed from 70% EtOH in to 100% EtOH and left in 100% EtOH overnight. The next day, hearts were incubated in 50-50 100%EtOH-Xylene at room temperature for 30 minutes, followed by two washes in 100% Xylene at 65 degrees, 1hr in 50-50 Xylene-Wax at 65 degrees C, two incubations in Wax at 65 degrees C for 30 minutes and a final 45 minute incubation in wax at 65 degrees C, before embedding in Wax. Wax sections were cut at 8um thick, with alternating sections split between three different groups for analysis of elastin, proteoglycans and collagen within the same heart.

For whole adult histology, embryos were cut to remove the trunk and washed from 70% EtOH into 100% EtOH and left in 100% EtOH overnight. The next day, hearts were incubated in Xylene at room temperature for 30 minutes twice, followed by 1hr in 50-50 Xylene-Wax at 65 degrees C and four incubations in Wax at 65 degrees C for 1hr, before embedding in Wax. Wax sections were cut at 8um thick and stained with Haematoxylin and Eosin (H&E) using standard protocols.

### ECM stains on paraffin sections

Wax sections were cleared in Xylene and rehydrated to MQ H_2_O. To stain for Elastin, slides were incubated in Miller’s Elastin (VWR, 351154S) for 2hrs (mouse) or 1hr. (zebrafish), rinsed in MQ, incubated in fresh 3% Iron (III) Chloride for 20 mins. (mouse) or 10mins (zebrafish) (Merck, 157740), rinsed in running tap water and counterstained in 10% Eosin Y (Thermo, 6766010) for 5 minutes. Alcian Blue staining (abcam AB150662) was carried out according to manufacturer’s instructions. Prior to collagen staining, rehydrated sections were incubated in Bouin’s solution (Sigma HT10132) for 16hrs (mouse) or 20hrs (zebrafish) overnight at room temperature washed under running tap water for 8 mins., rinsed for 2mins in ddH_2_O before carrying out Masson’s Trichrome staining according to manufacturer’s instructions (Sigma H15-1KT, HT1079). Once stained, all slides were cleared in Xylene, dehydrated and mounted in Histomount (National Diagnostics HS-103).

### Immunohistochemistry

Embryos were rehydrated from 100% MeOH to PBST and rinsed in PBS-X (1xPBS with 0.2% Triton-X (Sigma T8787) and then blocked for 1 hr at room temperature in 5% Normal Goat Serum (Fisher Scientific 10098792), 10mg/mL Bovine Serum Albumin (A2153, Sigma) in PBS-X (blocking buffer). The following primary antibodies were used: Cardiac Troponin (DSHB CT3, Mouse IgG2a, 1:200), Tagln (Stratech GTX125994, Rabbit, 1:1000), BrdU (Abcam ab6326, Rat, 1:200), GFP (Abcam 13970, Chick, only used for BrdU and FISH, 1:500), DsRed (also binds mCherry, Takara Bio 632496, Rabbit, 1:200), Isl1/2 (DSHB 39.4D5, Mouse IgG2b, DSHB 1:50), MLCK (Merck SAB4200808 (clone K36), Mouse IgG2b, 1:400), Sox9 (abcam ab185230, Rabbit, 1:500). Primary antibodies were incubated in blocking buffer and 1% DMSO (Thermo, 042780-AK) at 4C overnight with agitation. The next day, embryos were rinsed in PBS-X and washed thoroughly over two hours at room temperature. The following secondary antibodies were used (Thermo): Donkey anti-Mouse 647 (A31571), Goat anti-Rabbit 594 (A11012), Donkey anti-Rat 594 (A21209), Goat anti-Chick 488 (A11039), Donkey anti-Rabbit 568 (A11042), Goat anti-Mouse IgG2a 350 (A21130), Goat anti-Mouse IgG2b 594 (A22145), Donkey anti-Rabbit 647 (A31573), Goat anti-Mouse IgG2a 647 (A21241), Goat anti-Mouse IgG2a 488 (A21131) and Goat anti-Mouse IgG2b 647 (A21242). Secondary antibodies were incubated in blocking buffer at 1:200 dilution with 1% DMSO at 4C overnight with agitation. The next day, embryos were rinsed three times in PBS-X and rapidly dehydrated to 100% EtOH for resin embedding, which is described in the supplementary methods.

### Generation of *elnb* probe

The *elnb* (ENSDARG00000069017) probe was cloned using primers Forward: 5’-ATTAGGGGCTGGTGTTGGAA-3’ and Reverse: 5’-CCAAGTCCAAATCCAGCACC-3’ from 76hpf WT (AB) cDNA. The 1089bp fragment was ligated into pCRII-TOPO (Fisher 11503837) and sequenced by Sanger sequencing (Eurofins) to confirm insertion. For antisense mRNA, the plasmid was linearised with BamHI (NEB R0136) and transcribed using T7 (Promega, P207E) in the presence of Dioxygen-labelled nucleotides (Merck, 11277073910). mRNA in situ hybridisation protocols are listed in the supplemental methods.

### Miller’s Elastin staining of embryonic zebrafish

Embryos were rehydrated to MQ and incubated in Miller’s Elastin for 8hrs at room temperature, rinsed in MQ, incubated in freshly made 3% Iron (III) Chloride for 1hr at room temperature, rinsed in MQ and incubated in Eosin Y overnight for 15hrs at room temperature. The next day, embryos were rinsed in MQ and dehydrated to 100% EtOH for resin embedding.

### 3D reconstruction

Serial sections of resin embedded adult zebrafish hearts, cut at 7μm thick were stained with Haematoxylin and Eosin, photographed, and reconstructed using Amira 2020.2 (Thermo).

### Study design and statistical analysis

Each experiment was performed at least twice, composing of embryos from different clutches from different parents collected on different days. BrdU analysis was performed three times, with each experimental replicate represented by five embryos, in the same embryos, number of cells was quantified in the primordia. Sample size was pre-determined before analysis. For all other analyses individual embryos represent an experimental replicate and sample size was not pre-determined. H&E and Miller’s Elastin on embryos (<5dpf) was performed on both WT(AB) and Casper *Tg(kdrl:GFP)* genotypes. No differences were observed. All statistical analyses were carried out in GraphPad Prism.

## Results

### Conservation of arterial valve structure in adult zebrafish

We have previously characterised the morphology of the arterial roots in mice and human^7,17,34^. For comparison, we extended descriptions of the adult zebrafish arterial valve^25,26,35^, by three-dimensional reconstruction (Fig. 1A-B). As in amniotes, the zebrafish arterial valve is hinged at the distal extreme of the ventricle with the leaflets extending beyond the myocardial-smooth muscle boundary and have a plane of opposition along the anterior-posterior axis of the embryo (Fig. 1B-E, Fig S1). We also sectioned whole adults along the anterior-posterior axis and could clearly identify two leaflets that were aligned almost perpendicular to the left-right axis of the embryo (Fig. S1). To avoid confusion with mouse/human arterial leaflets, we have defined these as the dextral and sinistral leaflets given their predominant anatomical position within the adult (Fig. 1B, Fig S1). Overall, the two leaflets had no distinguishing morphological features and there was no continuity between either leaflet and the atrioventricular valve (Fig. 1B-C). Sections cut across the short axis of the outflow clearly identified sinuses between the leaflet and the wall (Fig. 1A, D-E). The tips of the leaflets were particularly thin and difficult to visualise in sections (Fig. 1C), but evaluation in multiple planes suggested a broad zone of apposition at their free edges (Fig. 1B-E).

**Figure 1.**
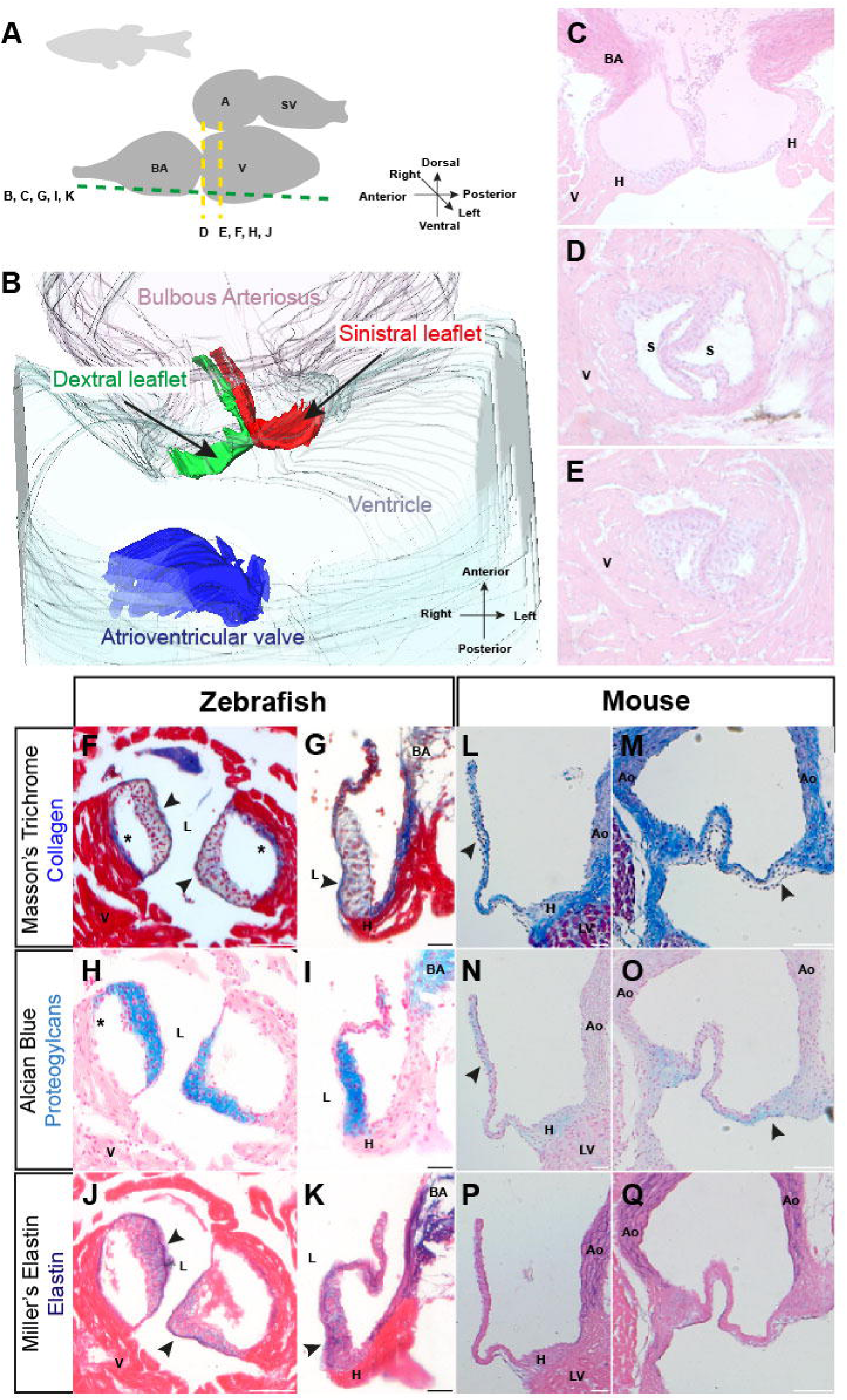
The zebrafish arterial valve is anatomically similar to other vertebrate arterial valves. **(A)** Overview of the orientation of the heart in adult zebrafish and plane of imaging in B and sections for C-K. **(B)** 3D reconstruction of the arterial pole, viewed ventrally. The arterial valve leaflets (green, red) are hinged in the ventricular myocardium and span the myocardial (grey)-arterial (pink) boundary and are not in continuity with the atrioventricular valve (blue). The arterial valve leaflets are aligned along the anterior-posterior axis of the zebrafish and are defined by their position within the adult: dextral (green) and sinistral (red). **(C)** Haematoxylin and Eosin stained long axis resin sections of excised adult zebrafish heart (Female, 9 months old) (n=4). **(D-E)** Haematoxylin and Eosin stained short axis resin sections of excised adult zebrafish heart (Male, 9 months old) at the level of distal (D) and proximal myocardium (E) showing two leaflets of the arterial valve, with sinuses between the leaflets and the wall (n=4). **(F)** Masson’s Trichome stained short axis wax section of excised adult zebrafish heart (Female, 14 months) at the proximal myocardium. Collagen (blue) is present in the lumen facing portion of the leaflets (arrowheads) and the wall of the sinus (asterisks) but is excluded from the interstitium. **(G)** Masson’s Trichrome stained long axis wax section of excised adult zebrafish heart (Female, 12 months old) showing the sinistral leaflet. Collagen is absent from the middle of the valve leaflet, with enrichment at the luminal surface (arrowhead), the root and the Bulbous Arteriosus (BA). **(H)** Alcian Blue stained short axis wax sections of same heart shown in (F). Proteoglycans (blue) are present throughout the interstitium of the leaflets, with very weak signal in the sinus wall (asterisk). **(I)** Alcian Blue stained long axis wax section of same heart shown in (G), showing the BA is rich in proteoglycans. **(J)** Miller’s Elastin stained short wax axis sections of same heart shown in (F, H). Elastin (purple) is present throughout the leaflet with clear enrichment of elastin on the luminal surface of the leaflets and in the sinus wall (arrowheads). **(K)** Miller’s Elastin stained long axis wax section of the same heart shown in (G, I) Elastin is present in the BA, enriched in the lumen facing aspect of the arterial leaflets and present in the root of the valve. (For A, C, E: n = 2, for B, D, F: n =4). **(L-M)** Masson’s Trichrome stained long axis section of Postnatal day (P) 90 mouse aortic valve, showing left (L) and non-coronary leaflet (M). Collagen is present in the aortic root hinge and wall. In the leaflets, Collagen is mainly localised to the arterial aspect and absent from the ventricular facing surface (L, M arrowheads). **(N-O)** Alcian blue stained long axis section same heart shown in (L-M). Sulfated proteoglycans are present in the aortic root hinge but largely absent for the wall of the aorta. In the leaflets, staining is reciprocal Collagen (N, O arrowheads). **(P-Q)** Alcian blue stained long axis section same heart shown in (L-O). Mature Elastin fibres are present in the wall of the aorta, with diffuse staining in the hinge. There are no clear fibres of elastin in the leaflets, diffuse signal is observed in the tips of the leaflets. C, G, I, K, L-Q: Anterior up, left: right. D, E, F, H, J: Dorsal: up, left: right. V: ventricle, BA: Bulbous Arteriosus, A: Atrium, SV: Sinus Venosus, LV: left ventricle, Ao: Aorta, S: Sinus, H: Hinge, L: Lumen. Scale bars: C-F, H, J, L, N, P: 50μm. G, I, K: 20μm. M, O, Q: 100μm.

In adult humans, it is widely described that the arterial valve leaflets have a trilaminar arrangement of extracellular matrix, with a fibrous layer of Collagen on their arterial aspect, a proteoglycan-rich middle layer, and an Elastin-rich layer on the ventricular side^9,10^. With this in mind, we investigated the localisation of these ECM components in the adult zebrafish arterial valve. Masson’s trichome staining revealed low levels of Collagen deposition in the leaflet interstitium of the adult arterial valve, with stronger staining on the luminal facing surface and sinus wall that overlies ventricular myocardium (Fig. 1F-G). Higher levels of staining was also found in the sinus walls and hinges (Fig. 1F-G). The BA was also strongly stained blue by Masson’s trichrome (Fig. 1G, S2A). In contrast, the whole of the leaflet interstitium stained strongly with Alcian blue (Fig. 1H-I), similar to human^9^ indicating an abundance of sulphated proteoglycans; this was largely restricted to the leaflets, although very low levels were found in the sinus wall (Fig. 1H, S2B), whilst the BA stained strongly (Fig. 1I, S2B). Mature elastin, revealed by Miller’s staining, was also found throughout the leaflets, with stronger staining on the ventricular side (Fig. 1J-K), as in adult human and overlapping Collagen (Fig. 2G). Elastin was also present in the sinus walls (Fig. 1J-K) although the hinges were deficient in elastin (Fig. 1K). The BA stained strongly for elastin, although we could not detect discrete elastin fibres (Fig. 1K, S2C). In summary, there is some evidence of leaflet lamination, with both collagen and elastin more abundant on the ventricular side, similar to what has been observed in human^9^.

**Figure 2.**
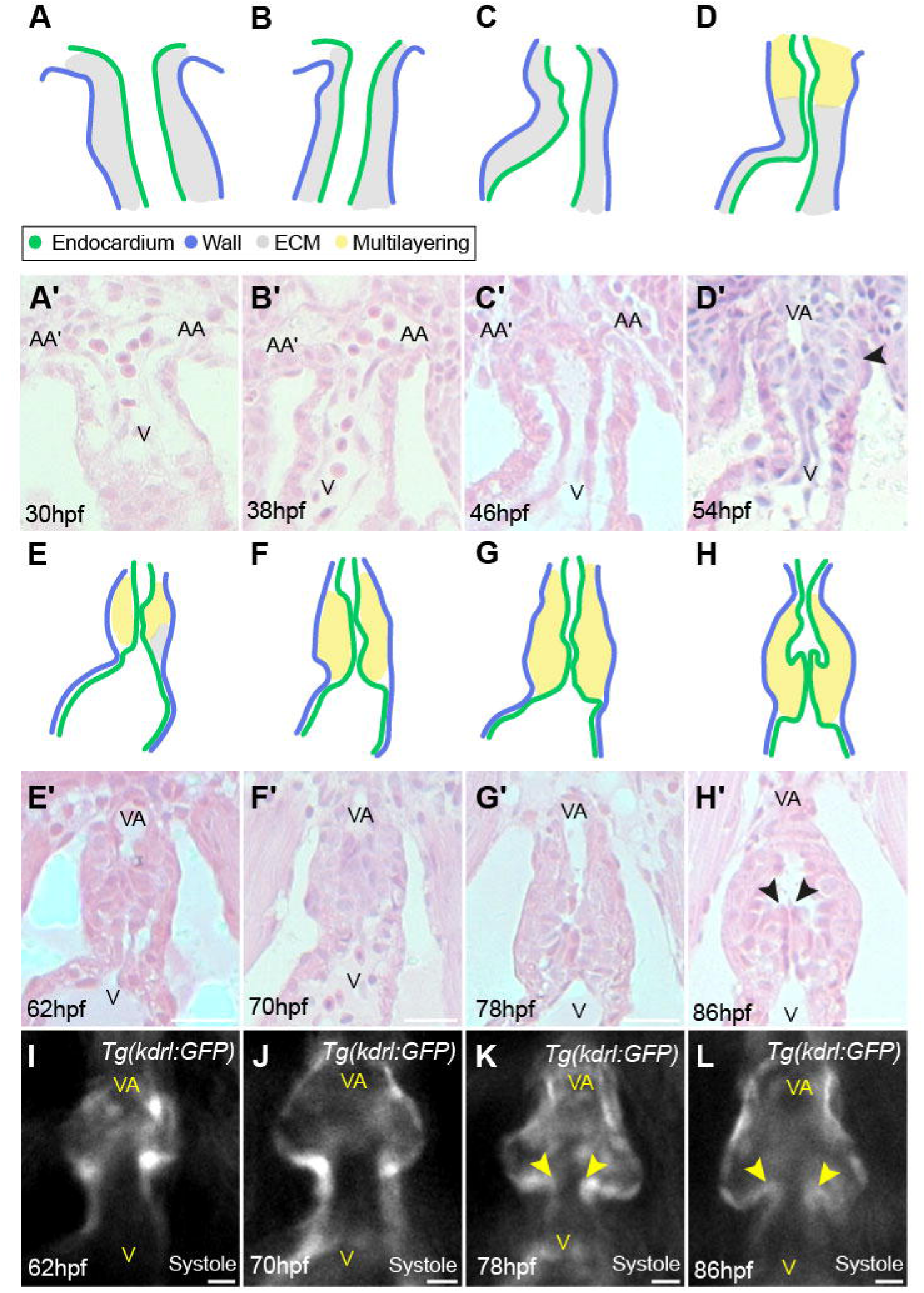
Development of the zebrafish arterial valve follows conserved events. **(A)** Schematic of 30hpf outflow tract (OFT). Two cell layers, the wall (lilac) and endocardium (green) are separated by the cardiac jelly (grey). **(A’)** Representative Haematoxylin and Eosin-stained midline resin section along the long axis of the OFT at 30hpf (n=6). **(B-C’)** Between 38-46hpf these two cell layers remain distinct and separated by ECM. (38hpf, n=7. 46hpf, n=10). **(D-D’)** At 54hpf, a multi-layering of cells is present at the distal-most point of the OFT where no ECM is visible (D’, arrowhead), with cells of unknown fate in yellow (n=8). **(E-E’)** ECM in the ventricle and OFT is largely absent at 62hpf (n=11). **(F-G’)**. Between 70-78hpf, any remaining ECM is lost and the OFT lengthens and tapers where it connects to the Ventral Aorta (70hpf, n=5. 78hpf, n=10). **(H-H’)** Sinuses are visible by 86hpf, defining two leaflets in the OFT (arrowheads) (n=5/8). **(I-L)** Live fluorescent imaging of the OFT between 62-86hpf of *Tg(kdrl:GFP)* embryos, marking the endocardium. Two primordia are visible at 62hpf (I) (n=13) and 70hpf (J) (n=14). The beginning of sinus formation is detectable at 78hpf (K) (n=11/15), with tips of leaflets visible (arrowheads) and is mostly complete by 86hpf (L) (n=17). AA: Aortic arches, V: Ventricle, VA: Ventral Aorta. Scale bars A-H: 20μm, I-L: 5μm.

As most of the data about arterial valve formation has been gathered from studies using mouse, we compared the zebrafish arterial valve with mouse more closely, analysing arterial valves from a 90-day old mouse (P90; adult). In the P90 mouse aortic valve, Collagen was present in the wall, hinge (Fig. S2D-E) and restricted to the arterial-facing aspect in all three leaflets (Fig. 1L-M, Fig S2E). As in the zebrafish, Alcian blue staining was present in a reciprocal pattern to Collagen in the valve leaflets, mostly present in the tips and also the hinge (Fig 1N-O, S2H). In contrast to the zebrafish BA, the mouse Aorta stained very weakly for with Alcian Blue (Fig. S2E). Finally, we detected minimal elastin in the adult mouse aortic valve leaflets, with only weak signal that was sparse and localised to the tips of the leaflets (Fig. 1P, Fig. S2I), whilst the wall of the aorta was rich in elastic fibres (Fig. 1P-Q, Fig. S2H). We also detected melanocytes in the mouse aortic valve leaflets (Fig. S2E, G, I), but not in the zebrafish arterial valve leaflets (Fig. 1C-E).

Together these data show that whilst elastin content and collagen localisation within arterial leaflets differs between zebrafish and mouse, there are clear and fundamental hallmarks of arterial valves. The arterial root is fibrous, the leaflets themselves rich in sulfated proteoglycans that do not co-localise with Collagen and the arterial aspect of the outflow is rich in Elastin and Collagen. Taken together, these analyses indicate general structural similarity between the adult zebrafish arterial valve and the adult mouse aortic valve. We next set out to investigate the developmental origins of the leaflets.

### Embryonic development of the arterial valve in zebrafish follows conserved events

Arterial valve formation can be viewed as a sequence of key events; establishment of primordia, sinus formation and leaflet sculpting^8,17,34^. To identify if these steps also occur in zebrafish, we examined histological sections of the developing arterial pole of the embryonic heart.

Initially, at 30hpf, the embryonic OFT is a single outer layer of myocardium, with an inner layer of endocardium, separated by extracellular matrix (ECM; cardiac jelly, grey)^36^ (Fig. 2A-B’). At 46hpf, the outflow has elongated through SHF addition at the distal end^13,37^, with no evidence of any arterial valve structures (Fig. 2C-C’). By 54hpf, the clear outer layer-ECM-endocardial layering is still observed in the proximal OFT (Fig. 2D-D’). Distally, however, little or no ECM is visible (arrowhead, Fig. 2D’), and a multi-layered arrangement of cells is apparent (Fig. 2D, yellow), forming a bulge that narrows the lumen of the vessel and the distinction between cell types lost. At 62hpf, the bulge of cells in the distal outflow tract appears larger and the ECM in the proximal OFT and the ventricle has thinned (Fig. 2E-E’). Between 70-78hpf, the OFT noticeably lengthens with a tapering connection to the ventral aorta (Fig. 2F-G’). At 86hpf, sinuses are visible between the bulge of cells and the outflow wall, for the first time delineating the two leaflets of the arterial valve. At the same time the OFT takes on a more pear-shaped profile around the forming valve (Fig. 2H-H’). Over the remaining course of embryonic development (86-118hpf), there is little change in the appearance of the arterial valve leaflets (Fig. S3A-D), although they appear thinner at 14dpf (Fig. S3E), indicating that sculpting happens during post-embryonic stages. We confirmed the window of primordia formation by quantifying the distance between the wall of the heart and the endocardium in *Tg(kdrl:GFP)* embryos in which all endothelial cells, including the endocardium express GFP (Fig. S3F). At 46hpf, wall thickness in the dOFT (distal outflow tract) was uniform (Fig. S3G-G’’), whilst at 54-70hpf we identified an increased wall thickness, coinciding with the multi-layered appearance of the dOFT (Fig. S3H-J’’). Thus, the bulge of cells that is seen in the distal OFT from 54hpf (Fig. 2D’) is the primordia of the arterial valve leaflets.

To extend and complement our histological analysis, we performed live imaging of the developing OFT to identify movement of valve leaflets at the earliest stages. In *Tg(kdrl:GFP)* embryos, the valve primordia could be visualised at 62-70hpf, but no leaflets were present (Fig. 2I-J, Movies 1-2). By 78hpf, rudimentary leaflets and sinuses could be distinguished, although the extent of their demarcation varied between embryos within the same clutch (Fig. 2K, Movie 3). By 86hpf, two leaflets, each with a distinct sinus, were identified in almost all embryos (Fig. 2L, Movie 4). In summary, the zebrafish OFT progress through the conserved, landmark events of arterial valve formation, from an initially open, continuous vessel (30hpf-46hpf), through primordia establishment (46hpf-70hpf), sinus formation (70hpf-86hpf) and finally leaflet sculpting (86hpf onwards).

### Arterial valve primordia form at the myocardial-arterial boundary of the outflow tract

To identify the position at which the arterial valve primordia form in zebrafish we performed immunohistochemistry for Cardiac Troponin (myocardium) and Transgelin (Tagln; formerly SM221Z), a well conserved marker of smooth muscle cells in the arterial pole^7,17,38^ in *Tg(kdrl:GFP)* embryos. At 46hpf, when there are no primordia present (Fig. 2, Fig. S3), the entire OFT wall is myocardial (Fig. 3A-A’). At 54hpf, the beginning of primordia formation, we could identify distally positioned cells that did not express Troponin nor Tagln (Fig. 3B-B’). By 62hpf, cells in the wall distal to the valve primordia were sparsely Tagln+ (Fig. 3C-C’) and by 70hpf, Tagln expression was robust in the dOFT. The arterial valve primordia were clearly seen within the myocardial portion of the OFT at the boundary of cardiomyocytes and smooth muscle cells and were negative for endocardial, myocardial, and smooth muscle markers (arrowheads, Fig 3D’). Thus, as in mouse and human, the arterial valve forms in conjunction with the arterial root at the myocardial-smooth muscle (myocardial-arterial) boundary and during this window, the length of the OFT increases (Fig. 3E).

**Figure 3.**
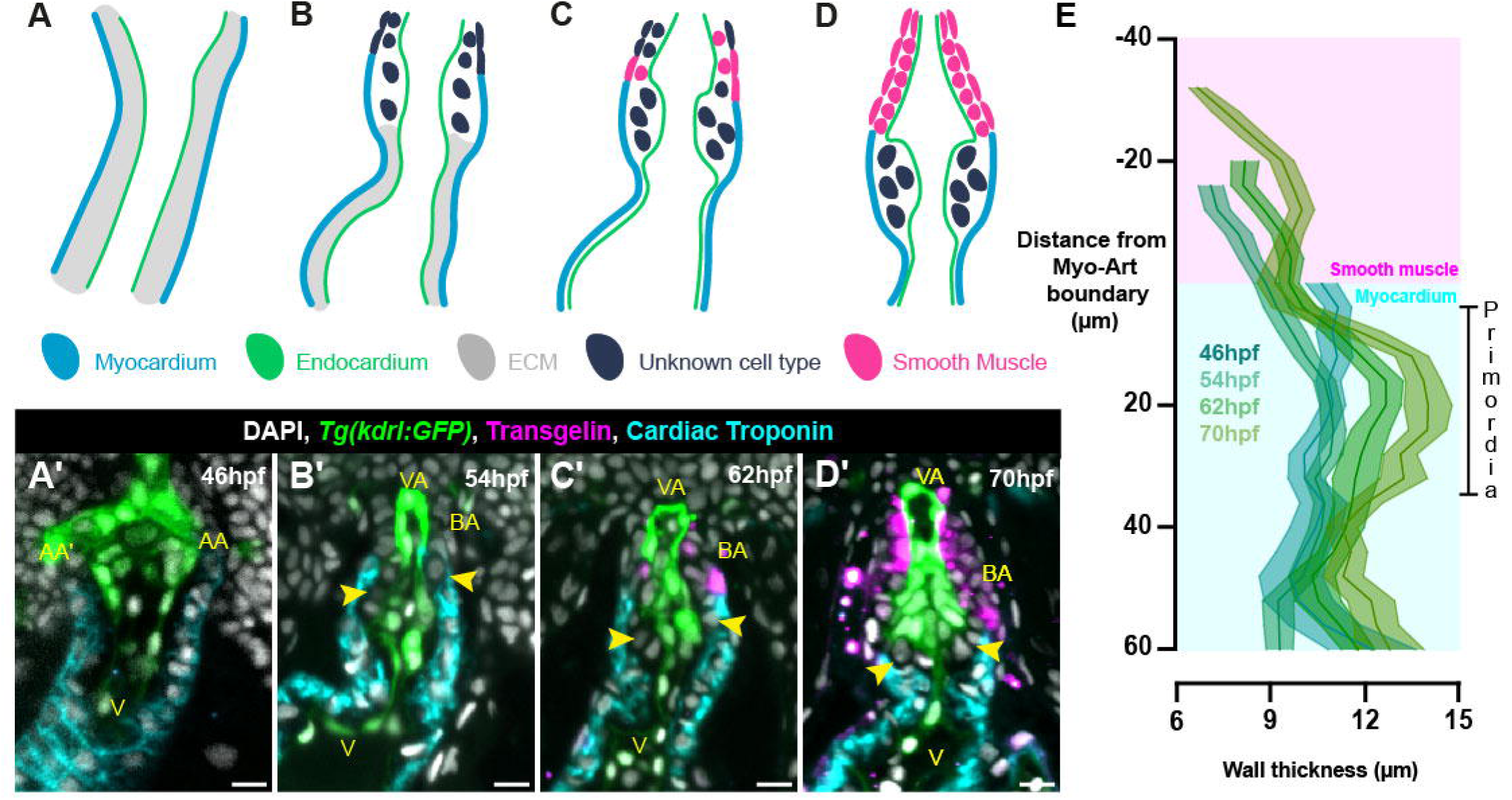
Arterial valve primordia form at the myocardial-smooth muscle boundary. **(A-D)** Schematics of OFT at 46hpf (A), 54hpf (B), 62hpf (C) and 70hpf (D). **(A’-D’)** Representative midline resin sections of *Tg(kdrl:GFP)* (endocardium, green) embryos stained for Tagln (smooth muscle, magenta), Cardiac Troponin (myocardium, cyan) and DAPI (white) between 46-70hpf. At 46hpf (A-A’) the entire OFT is myocardial (n=12), at 54hpf (B-B’) there is a region distally that is not myocardial and does not express Tagln (asterisk), some cells are identifiable between the endocardium and myocardium (arrowheads) (n=15). By 62hpf (C-C’), the region distal to the myocardium begins to express Tagln (n=9), which is more robust at 70hpf (D-D’) (n=11). Cells present between the endocardium and myocardium do not express any markers (C’, D’ arrowheads). **(E)** Quantification of wall thickness, averaged across left and right, with distance along OFT measured relative to myocardial-arterial boundary (46hpf n=24, 54hpf n=30, 62hpf n=18, 70hpf n=22). The primordia of the arterial valve form at the distal most point of the ventricular myocardium. E: Mean ± S.E.M. AA: Aortic arches, V: Ventricle, VA: Ventral Aorta, BA: Bulbous Arteriosus. Scale bars: 10μm.

A number of studies have characterised proliferation in the zebrafish OFT^39,40^, but not in the context of arterial valve development. We therefore evaluated proliferation using timed pulses of BrdU during primordia formation, defining cells of the primordia by their relative position to the myocardium and endocardium and absence of expression of these two markers (Fig. 3). Between 54-70hpf, cell number in the valve primordia increases (Fig. S4D-D’) and this increase is associated with cells undergoing division as demonstrated by incorporation of BrdU (Fig. S4E-E’). This occurred equally in the left and right primordia (Fig. S4D’, E’).

It has been previously suggested that cells of the valve primordia are smooth muscle, based mainly on analysis of Elastin localisation^28^. However, in our analysis, cells within the arterial valve primordia did not express the known smooth muscle marker Tagln (Fig. 3B’-D’). To clarify this, we examined the spatio-temporal expression of *elastin b* (*elnb),* by wholemount mRNA *in situ* hybridisation. This identified expression in the distal outflow from 54hpf onwards (Fig. S5A-D’) coinciding with primordia formation (Fig. 2, 3) and smooth muscle appearance in the BA (Fig. 3B-D’). However, midline sections in *Tg(kdrl:GFP)* embryos at 54 and 70hpf revealed no *elnb* expression in cells of the valve primordia (yellow region, Fig. S5F-G’’). Which was further confirmed by Miller’s elastin staining of midline sections at 46-70hpf (arrowheads, Fig. S5G-J). Therefore, our data demonstrate that together with smooth muscle, myocardium and endocardium, a fourth population of cells, are present in the developing OFT that give rise to the arterial valve leaflets.

### Arterial valve primordia form through direct differentiation of second heart field progenitors

The arterial valve primordia in mouse and human form through two distinct mechanisms: either via cellularisation of an ECM-rich cushions by EndoMT-derived cells and neural crest cells, or via direct differentiation from SHF progenitors in the ICVS^17,18^.

To establish the origin cells in the zebrafish arterial valve primordia, we used Cre-lox-mediated lineage tracing to identify the contribution of endothelial cells and neural crest cells. In *Tg(kdrl:Cre); Tg(ubi:loxP-EGFP-loxP-mCherry)* embryos at 70hpf, we observed recombination of the reporter in the endocardium (Fig. 4B-B’, magenta), identical to that of *Tg(kdrl:GFP)* (Fig. 2-3, S5), but no recombination in the cells of the arterial valve primordia (yellow region, Fig. 4B’). Conversely, at the same stage, the cells within the atrioventricular valve showed almost total recombination (Fig. 4C-C’)^32,41^. To establish the contribution of neural crest cells to the arterial valve, we generated a new allele of the previously reported *Tg(sox10:iCre,cryaa:DsRed2)*^42^ (Fig. S6). In *Tg(sox10:iCre, cryaa:DsRed2); Tg(ubi:loxP-EGFP-loxP-mCherry)* embryos at 70hpf, we could identify a neural crest cell contribution to the distal-most portion of the BA and ventral aorta^43^, but no recombination was seen in either valve primordia (yellow region, Fig. 4D-D’ and 4E-E’).

**Figure 4.**
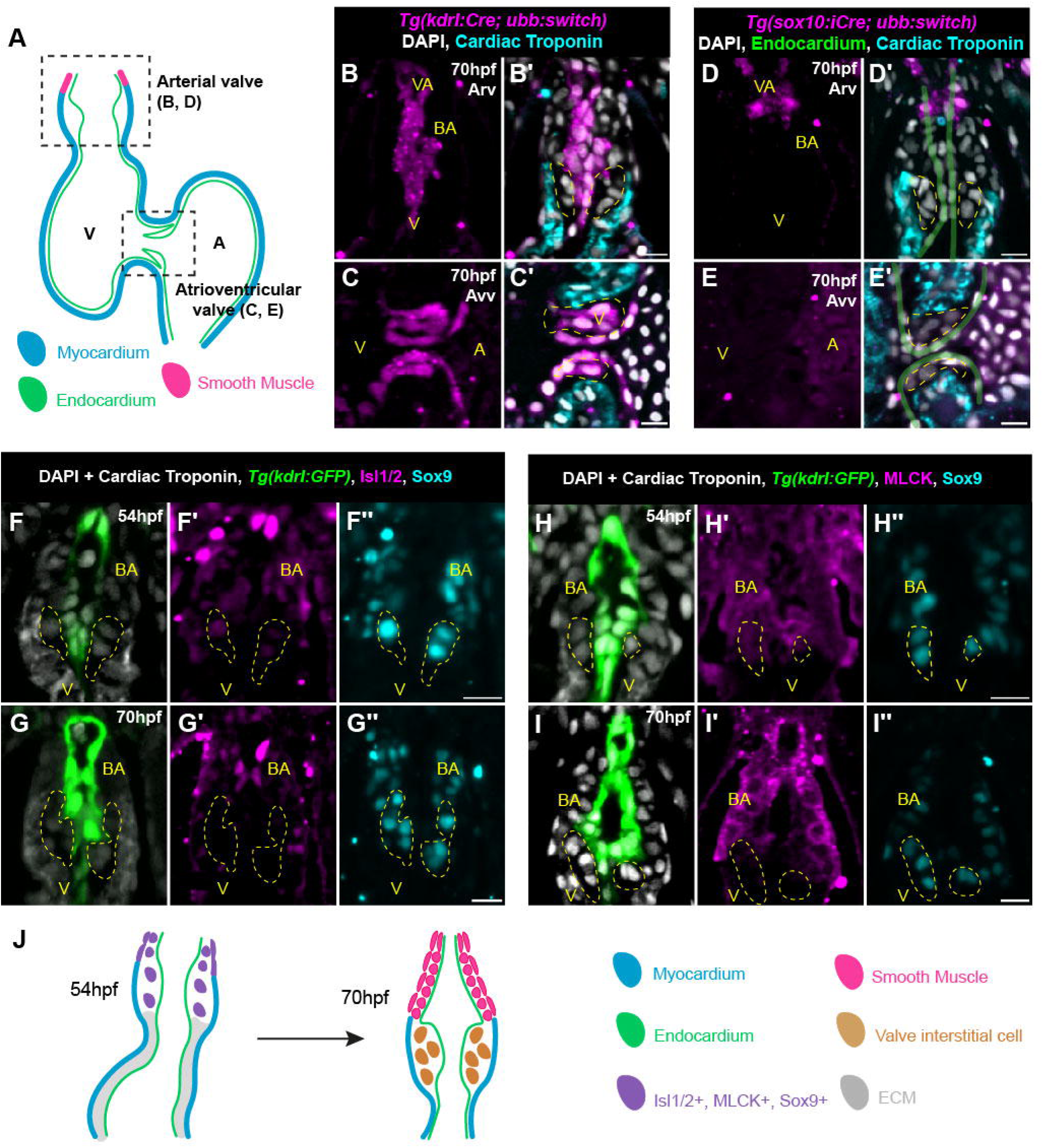
Direct differentiation of SHF progenitors establishes the zebrafish arterial valve. **(A)** Schematic of embryonic heart at 70hpf, showing positions of the two valves examined in Cre-lineage tracing analyses. **(B-C’)** Lineage tracing of endothelial cells at 70hpf in *Tg(kdrl:Cre); Tg(-3.5ubb:loxP-EGFP-loxP-mCherry)* embryo (magenta). (B’) Cells of the arterial valve primordia (yellow), identified between distal-most myocardium (cyan) and endocardium are not of endocardial origin (n=19). (C’) Cells of the Avv (yellow) are derived from the endocardium (n=19). **(D-E’)** Lineage tracing of neural crest cells at 70hpf in *Tg(sox10:iCre, cryaa:DsRed2); Tg(-3.5ubb:loxP-EGFP-loxP-mCherry)* embryo (magenta). (D’) Cells of the arterial valve primordia (yellow), identified between distal-most myocardium (cyan) and endocardium (green line) are not of neural crest origin. Recombination is present in the distal tip of the BA (n=8/10). (E’) Cells of the Avv (yellow) are not derived from the neural crest (n=12). **(F-F’’)** Representative midline resin section of immunohistochemistry on *Tg(kdrl:GFP)* (green) embryo at 54hpf for Cardiac Troponin (white, membrane), DAPI (white, nuclear) (F), Islet1/2 (Isl1/2, magenta) (F’) and Sox9 (cyan) (F’’). Cells of the distal OFT, including within the arterial valve primordia (yellow) co-express Isl1/2 and Sox9 (n=19). **(G-G’’)** Representative midline section of immunohistochemistry on *Tg(kdrl:GFP)* embryo at 70hpf. Cells within the arterial valve primordia (yellow) have downregulated Isl1/2 (G’) but maintain expression of Sox9 (G’’) defining them as valve interstitial cells (n=12). **(H-H’’)** Representative midline section of immunohistochemistry on *Tg(kdrl:GFP)* (green) embryo at 54hpf for Cardiac Troponin (white, membrane), DAPI (white, nuclear) (F), Myosin Light Chain Kinase (MLCK, magenta) (F’) and Sox9 (cyan) (F’’). Cells distal to the myocardium, including those in the arterial valve primordia (yellow) show diffuse MLCK expression and co-express Sox9 (n=19). **(I-I’’)** Representative resin midline section of immunohistochemistry on *Tg(kdrl:GFP)* embryo at 70hpf. MLCK expression in the BA is now restricted to the membrane and complementary to the Cardiac Troponin. Cells within the arterial valve primordia (yellow) have downregulated MLCK (G’), but maintain expression of Sox9 (G’’) (n=13). **(J)** Summary of expression data from (F-I). V: Ventricle, VA: Ventral Aorta, BA: Bulbous Arteriosus, Arv: arterial valve, Avv: atrioventricular valve. Scale bars: 10μm.

As these lineage tracing experiments demonstrate that the origin of the arterial valve primordia are EndoMT- and neural crest cell-independent, we asked whether there might be direct SHF differentiation, as seen in the ICVS of mouse and human^17,18^. As this mechanism is closely linked with the transition zone in the dOFT^15,17^, we first examined whether the transition zone was also present in the arterial pole of the zebrafish heart. As with other vertebrates, zebrafish SHF addition requires the conserved Islet family of transcription factors^13,44^, with Isl1+, Isl2a+ and Isl2b+ cells present in the arterial pole, but only *isl2b* required for cell addition^44^. At both 46hpf (Fig. S7A-D’’’) and 54hpf (Fig. S7E-H’’’), we could clearly identify cells at the arterial pole of the heart co-expressing Cardiac Troponin and Isl1/2, confirming the presence of the transition zone in the zebrafish heart.

We next examined the expression patterns of Isl1/2 and Sox9 in the arterial valve primordia as co-expression of these genes is the hallmark of the direct differentiation mechanism of primordia formation^17^. At 54hpf, we identified Isl1/2+ Sox9+ cells within the arterial valve primordia (yellow region, Fig. 4F-F’’) and distal to the forming valve in the outflow wall, that forms the BA. By 70hpf, the cells in the primordia had down-regulated Isl1/2 but maintained Sox9 expression (yellow region, Fig. 4G-G’’). Thus, the cells in the valve primordia are formed by direct differentiation of Isl-expressing SHF progenitors into VICs. At 70hpf, cells in the developing BA were still Isl1/2+ (Fig. 4G’), indicating late differentiation from SHF, but also co-expressed Sox9 (Fig. 4G’’). As this suggested that smooth muscle cells of the BA are Sox9+, we confirmed the smooth muscle phenotype of the developing BA at this stage using a second marker, Myosin Light Chain Kinase (MLCK)^45^. At 54hpf, we observed diffuse MLCK signal, complementary to Cardiac Troponin and overlapping with Sox9+ cells both in the root and smooth muscle population (yellow region, Fig. 4H-H’’). At 70hpf, MLCK expression distal to the myocardium was membrane restricted (compare Fig. 4H’ with 4I’), and overlapped with Sox9, confirming that the developing BA of the zebrafish expresses Sox9. Similar to Isl1/2 (Fig. 4F’, G’), we observed a reduction in MLCK signal in the Sox9+ cells of the arterial valve primordia between 54hpf and 70hpf (compare Fig. 4H’ to Fig 4I’). Taken together, the formation of the arterial valve primordia occurs at the transition zone, through direct differentiation of SHF progenitors into Sox9+ VICs (Fig. 4J), defining arterial valve establishment in zebrafish to be a distinct mechanism from that of atrioventricular valve development.

### Abnormal second heart field development results in defective arterial valve primordia development

Having characterised the progenitor populations required for development of the zebrafish arterial valve, we sought to confirm our findings in mutant lines that impact SHF development. The transcription factor Tbx1 has an evolutionarily conserved role in maintaining the SHF progenitor pool^46,47^ with loss of *TBX1* resulting in DiGeorge Syndrome^48^. *tbx1* mutant zebrafish have been shown to have a shorter OFT^27,37,46^, but the impact on the arterial valve remains unresolved.

At 70hpf, *tbx1* mutants have fully penetrant pericardial odema, absence of pharyngeal arches, and an abnormally positioned OFT (Fig. 5A-B’). Using *elnb* to mark the BA revealed that this structure is either severely hypoplastic (Fig. 5D-D’) or absent (Fig. 5D’’) in *tbx1* mutants. Whilst WT siblings have a regular, pear-shaped aortic root housing the arterial valve primordia (Fig. 5E), *tbx1* mutants have a grossly disorganised OFT (Fig. 5F). Expression of MLCK in WT siblings was robust and membrane restricted, enabling VICs to be defined by their absence of myocardial, endocardial or smooth muscle marker (yellow region, Fig. 5E). In *tbx1* mutants, MLCK expression was often totally absent from the OFT, showing variable expression between individual embryos, likely reflecting the spectrum of OFT hypoplasia identified by analysis of *elnb* expression (Fig 5D-D’’). This prevented identification of VICs by absence of MLCK expression, therefore, as before, we defined VICs to be within the myocardial collar, under the endocardium, but negative for both of these markers (Fig. 3, S4). This approach identified is a significant reduction in number of VICs, with both primordia impacted similarly (Fig. 4G-G’), confirming that *tbx1* is required for both proper OFT development, and formation of the arterial valve primordia.

**Figure 5.**
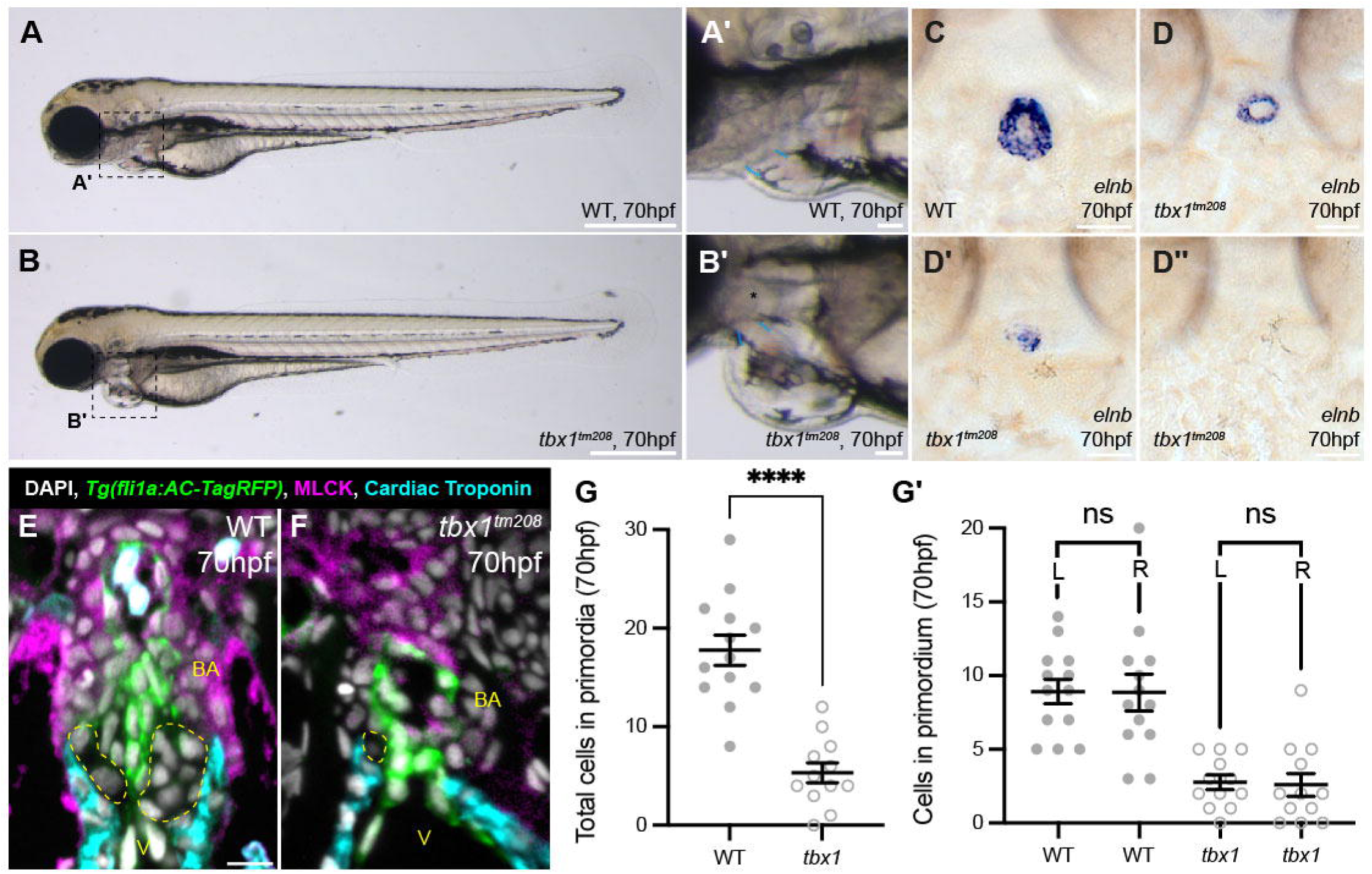
*tbx1* mutants have a dysmorphic OFT and hypocellular arterial valve primordia. **(A-B’)** Representative brightfield image of WT sibling (A, n=6) and *tbx1^tm208^* homozygous mutant (B, n=12) at 70hpf, dashed boxes in (A, B) highlights heart shown in (A’, B’). In *tbx1^tm208^* mutants (B’), the pharyngeal arches are absent (asterisk), and the angle of the OFT angle is steeper than in WT (blue). **(C-D’’)** Representative images of mRNA *in situ* hybridisation for *elnb* in WT sibling (C) and *tbx1^tm208^* homozygous mutants (D-D’’) at 70hpf. tbx1 mutants display variability in size of *elnb* domain. **(E-F)** Representative midline sections of immunohistochemistry on WT sibling (E) and *tbx1^tm208^* homozygous mutants (F) carrying *Tg(fli1a:AC-TagRFP)* to mark the endocardium (green), cardiac troponin (cyan) and MLCK (magenta). The *tbx1^tm208^* OFT is smaller and dysmorphic with diffuse MLCK expression. **(G-G’)** Quantification of number of cells in arterial valve primordia (yellow) at 70hpf in WT sibling and *tbx1^tm208^* homozygous mutants at 70hpf. Loss of *tbx1* results in a significant reduction in number of valve interstitial cells, and impacts the left and right primordia equally. (WT sibling, n=13. *tbx1^tm208^*, n=12). G-G’: Mean ± S.E.M, Welch’s unpaired t-tests. ****: p<0.0001, ns: not significant. V: Ventricle, BA: Bulbous Arteriosus. Scale bars: A, B: 50μm, A’, B’: 50μm, C-D’: 20μm, E-F: 10μm.

We have previously shown that SHF-specific ablation of core PCP component *Vangl2* in mice disrupts organisation of the transition zone, resulting in smaller or misplaced ICVS, dysplastic aortic leaflets and BAV^15,17^. As the OFT phenotype of *vangl2* mutant zebrafish has never been characterised, we investigated possible conservation of function. At 70hpf, the *vangl2* mutant zebrafish has a shorter antero-posterior axis (Fig. 6A, B), but overtly, heart morphology appears normal (Fig. 6A’, B’). Analysis of *elnb* expression identified that the BA is significantly larger, but less round in *vangl2* mutants compared to siblings at 70hpf (Fig. 6C-F). MLCK expression in *vangl2* mutants was comparable to WT siblings (Fig. 6G-H) and its membrane restriction suggested proper maturation of smooth muscle cells in *vangl2* mutants (Fig. 4H-I’’). Despite the abnormal shape of the OFT in *vangl2* mutants (Fig. 6C-H), the number of VIC in the arterial valve primordia was not different to WT siblings (Fig. 6I-I’). Together, these changes suggest that loss of *vangl2* leads to misshapen arterial pole, similar to the appearance in the mouse *Vangl2* mutant^15^.

**Figure 6.**
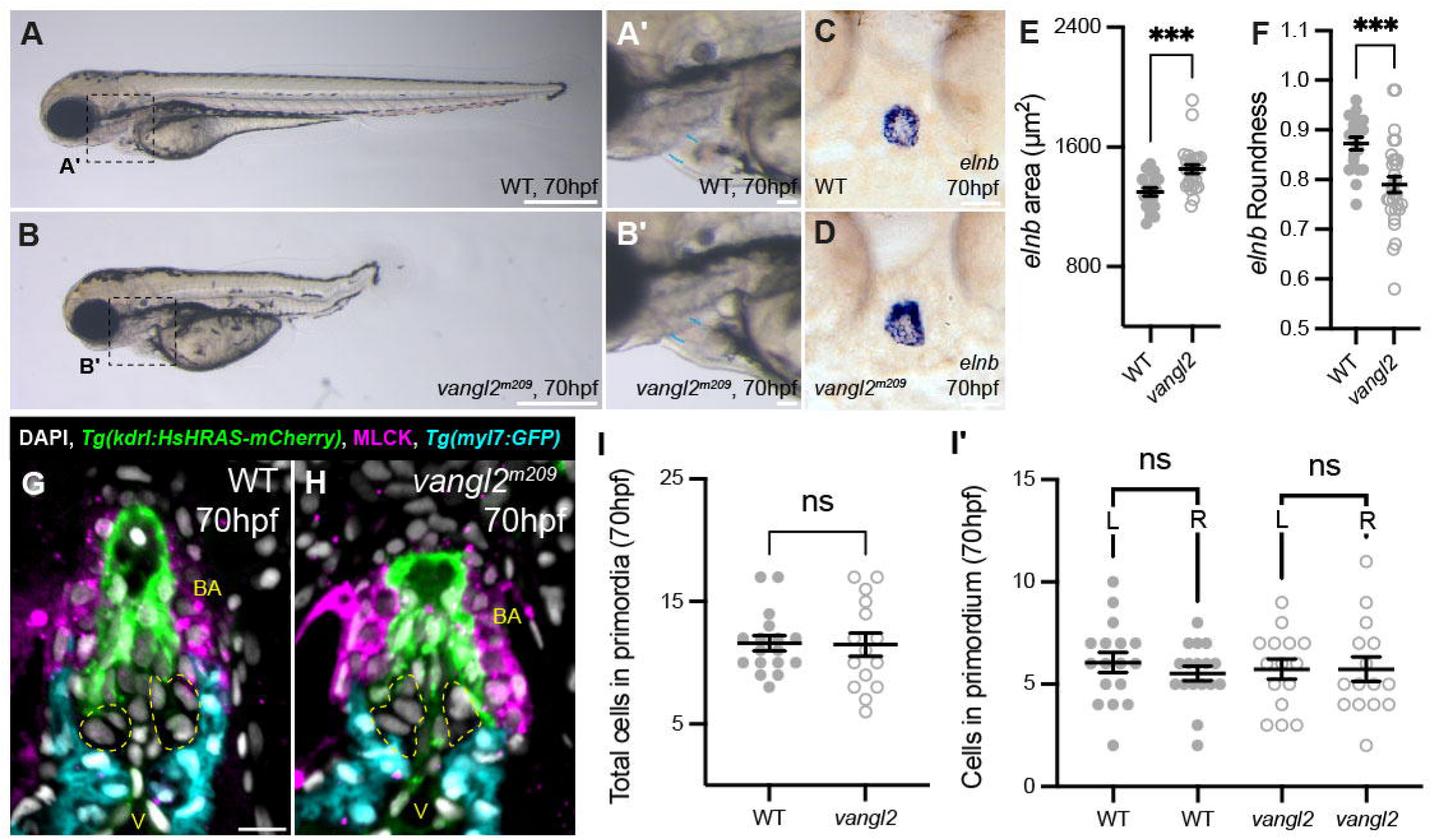
*vangl2* mutants have misshapen OFT. **(A-B’)** Representative brightfield image of WT sibling (A, n=7) and *vangl2^m209^* homozygous mutant (B, n=15) at 70hpf, dashed boxes in (A, B) highlights heart shown in (A’, B’). The overt morphology of the *vangl2^m209^* OFT appears normal (blue). **(C-D)** Representative images of mRNA *in situ* hybridisation for *elnb* in sibling (C) and *vangl2^m209^* homozygous mutant (D) at 70hpf. **(E)** Quantification of area of *elnb* domain at 70hpf. Loss of *vangl2* results in a larger elnb domain. **(F)** Quantification of roundness of *elnb* domain at 70hpf. Loss of vangl2 leads to a less round *elnb* domain (Siblings, n=19. *vangl2^m209^*, n=28). **(G-H)** Representative midline sections of immunohistochemistry on WT sibling (G) and *vangl2^m209^* homozygous mutants (H) carrying *Tg(kdrl:HsHRAS-mCherry)* to mark the endocardium (green) and *Tg(myl7:GFP)* to mark the myocardium (cyan) and MLCK (magenta). The *vangl2* OFT appears squat, with the myocardial collar less pronounced. **(I-I’)** Quantification of number of cells in arterial valve primordia (yellow) at 70hpf in WT sibling and *vangl2^m209^* homozygous mutants at 70hpf. Loss of *vangl2* does not impact number of valve interstitial cells (WT sibling n=17. *vangl2^m209^*, n=15). E, F, I-I’: Mean ± S.E.M, Welch’s unpaired t-tests. ***: p<0.001, ns: not significant. V: Ventricle, BA: Bulbous Arteriosus. Scale bars: A, B: 50μm, A’, B’: 50μm, C-D: 20μm, G-H: 10μm.

In summary, we have shown that loss of *tbx1*, a known regulator of SHF development^27,46^, leads to a small, dysmorphic OFT and is accompanied by a dramatic loss of cells in the arterial valve primordia, supporting our data that these primordia are SHF-derived. Loss of *vangl2*, a gene known to be required for SHF deployment in mice^15^, leads to a misshapen OFT suggesting a conserved role for *vangl2* in arterial pole development.

## Discussion

In this study, we have characterised the mature structure and embryonic origins of the zebrafish arterial valve. We have shown that both leaflets are established through direct differentiation of SHF progenitors (Fig. 4, 7), which is comparable to the development of the anterior and non-coronary leaflets of the mouse pulmonary and aortic valves respectively^17^. However, it is important to appreciate that this is not a pulmonary, nor aortic valve as it has been termed in some studies.

There are clear differences in the overall appearance of the arterial valve in zebrafish and mammals. Most obviously, the zebrafish arterial valve is bicuspid (Fig. 1, S1-2) with no interleaflet triangles nor continuity between the arterial and atrioventricular valve (Fig. 1). There are valve sinuses, but no coronary ostia as the coronary circulation in fish originates after the gills. Despite these differences, a basal ring, concomitant with the ventriculo-arterial junction, can be seen and the distal attachments of the leaflets define a sino-tubular junction (Fig. 1). Histological analysis of ECM composition of the valve revealed structures that were comparable to those found in the adult mouse (Fig. 1, S2) and in humans. Enriched elastin deposition on the ventricular face of the arterial leaflets is in keeping with the ventricularis, whilst the spongiosa appears present marked by a proteoglycan-rich extracellular matrix. The fibrosa in zebrafish is not well established with only minimal collagen deposition which is present on the luminal aspect of the leaflet. Despite this, we could clearly identify a general, conserved, reciprocal localisation pattern of proteoglycans and collagen in the valve leaflets (Fig. 1, S2).

Both arterial valve primordia arise directly from undifferentiated SHF at the most distal extent of the OFT transition zone, with no contribution from neural crest cells or EndoMT (Fig. 4, 7, S7). As in the mouse, we observed no trans-differentiation from myocardium and ruled out a smooth muscle identity of these cells (Fig. S5). The conservation of direct differentiation of VICs from SHF progenitors, from fish to mammals, suggests a relation to the ancestral mechanism of arterial valve development, predating the co-option of neural crest for complete OFT septation. The presence of the transition zone in zebrafish (Fig. S7), chick^49^, mouse^15,17^, human^14^ and probably Xenopus^50^, a domain that is tightly linked with the site of valve primordia formation, supports this further. In Xenopus, the neural crest are not required for incomplete septation and remains in the aortic sac and its associated arches^51^. This is very similar to what we and others have observed in zebrafish, where they form part of the ventral aorta (Fig. 4)^43^. Altogether, this suggests that the OFT of zebrafish is composed of SHF-derived myocardium proximally, above this, SHF-derived smooth muscle, and most distally, neural crest-derived smooth muscle. These contributions from SHF and neural crest cells, again show a high level of conservation with the mouse OFT; SHF generates the smooth muscle at the base of the aorta and pulmonary trunk, whilst the smooth muscle of the ascending aorta and associated arteries are of neural crest origin^7,52^. Interestingly, we observed strong Sox9 signal present in the smooth muscle of the OFT and other structures in the pharyngeal region (Fig. 4 and data not shown). In P1 mice, phosphorylated Sox9 is present in the wall of the root, supporting that Sox9 expression in the OFT wall is in fact a conserved pattern^22^.

**Figure 7.**
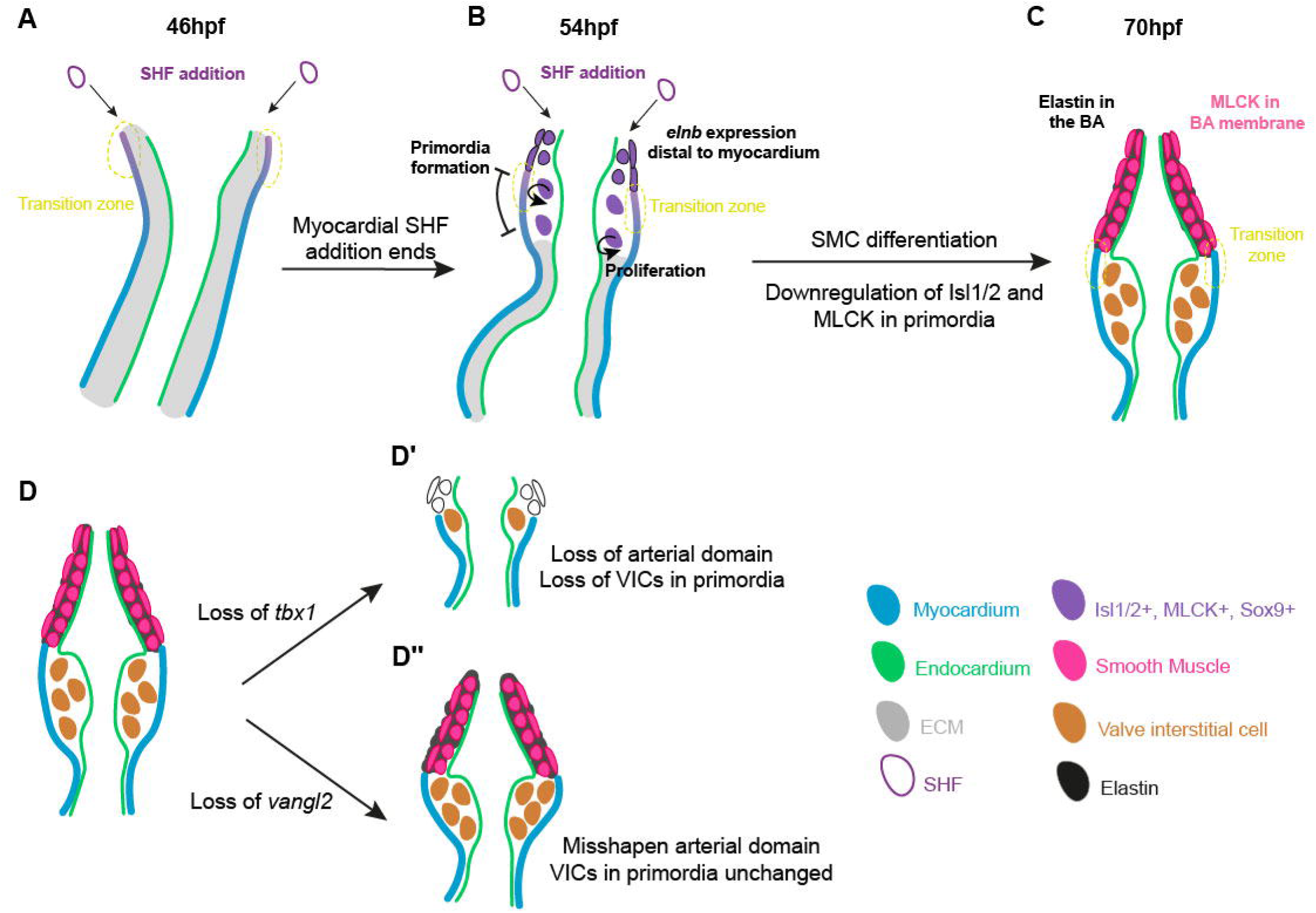
The zebrafish arterial valve forms by direct differentiation of SHF progenitors. **(A)** At 46hpf, addition to the OFT from the second heart field (SHF) occurs at the transition zone (yellow), where cells co-express Isl1/2 and mature cardiomyocyte markers, before downregulating Isl1/2. **(B)** At 54hpf, SHF addition no longer adds myocardium to the OFT, but instead smooth muscle which expresses *elnb*. This switch is the site of formation of the primordia at the transition zone, forming a bulge between the myocardium and endocardium. Cells within the early primordia proliferate and share similar expression patterns to the distal smooth muscle, but do not express *elnb*. **(C)** By 70hpf, the smooth muscle cells of the Bulbous Arteriosus (BA) are now more molecularly distinct from the cells of the arterial valve primordia. The smooth muscle expresses *elnb*, MLCK, Tagln and Sox9 and is surrounded by elastin fibres, whilst the cells of the primordia have downregulated Isl1/2 and MLCK but maintain Sox9. The primordia is devoid of cells expressing *elnb* or any elastin fibres. These are the interstitial cells of the arterial valve primordia. **(D)** Impact on arterial pole in mutants with improper SHF development. **(D’)** Loss of *tbx1*, results in a smaller, dysmorphic OFT with fewer VICs in the arterial valve primordia. The arterial domain is consistently smaller but is highly variable in size, most embryos have some expression of *elnb*, but total absence of MLCK expression, suggesting either a delay or loss in smooth muscle differentiation. **(D’’)** In *vangl2* mutants, the OFT appears to development normally, but is misshapen. There is a well-defined arterial domain and no impact on VICs in the arterial valve primordia.

The analysis of *vangl2* and *tbx1* zebrafish mutants provides important links between development of the zebrafish and mammalian outflow tracts (Fig. 7D-D’’). Loss of *Tbx1* in mice results in a shortened distal OFT with development of the anterior ICVS impacted^17,53^ and we have identified comparable phenotypes in the zebrafish *tbx1* mutant (Fig. 6, 7D’), mirroring the congenital heart malformations seen in DiGeorge syndrome. However, whilst the anterior ICVS is affected in *Tbx1* mice mutants due to its requirement for posterior SHF addition^53,54^, both arterial leaflets were impacted similarly in the zebrafish *tbx1* mutant (Fig. 5), suggesting that the unseptated OFT may not have the distinct regional identities observed in mouse. In contrast, loss of *vangl2* resulted in a more subtle defect, with sufficient SHF derived cells present in the primordia, but the overall size and shape of the smooth muscle domain marked by *elnb* was abnormal (Fig. 6). This suggests that similar to mouse^15^, loss of PCP signalling may lead to disordered cell addition to the arterial pole (Fig. 7D’’), rather than loss as observed in *tbx1* mutants. Little is known about human *VANGL2* mutations as these are likely to be incompatible with development^55^.

Disrupted aortic valve morphology in patients associated with altered haemodynamics and aortopathies, yet which abnormality (flow, valve structure or wall integrity) is the driver of the pathology remains unanswered^56–58^. Zebrafish are uniquely placed to investigate these interactions, due to high conservation of flow-dependent processes and accessibility for repeated *in vivo* live imaging. Previous studies have shown that cardiac function is required for the growth of the OFT^29^ ^39^ and *tnnt2a* morphants (which lack a heartbeat) are reported to have a complete absence of arterial valve primordia^28^. Additionally, it has been recently shown that disrupted TGF-β signalling in zebrafish result in dilatations of the arterial pole that are transcriptionally similar to those of Marfan Syndrome and Thoracic Aortic Aneurysm Dissection^59^. This, together with our description of the overt conservation of OFT structure, suggests the possibility of zebrafish as a model to investigate a wider repertoire of arterial pole malformations.

Altogether, this demonstrates a very high level of conservation of origin, expression profiles and gene function of the cells that reside in the developing OFT of the zebrafish heart. With the comparable development of the arterial valve to that of the anterior and non-coronary leaflets of the human arterial valves, this demonstrates that modelling OFT anomalies in zebrafish is possible and can provide clinical relevance. In particular, our mutant studies exemplify a range of outflow tract phenotypes that reflect the utility of zebrafish in variant analysis, most obviously subtypes of BAV where the SHF-derived non-coronary leaflet is hypoplastic or missing. Zebrafish are also likely to be useful in assessing variants in Tetralogy of Fallot - often found in DiGeorge Syndrome^60^ where all the affected structures originate from the SHF and the pulmonary valve is bicuspid, with at least 60% of these cases missing the SHF-derived anterior leaflet^6^.

There are obvious limitations to the use of zebrafish for BAV variant testing. The basis of some forms of BAV may be related to disruption of cardiac neural crest cell migration and outflow tract septation; these cannot be modelled in zebrafish. However, these events may be secondary to earlier defects in SHF addition, which then impacts neural crest cell migration^30,61^. Alternatively, it may be possible to use the EndoMT-driven process of atrioventricular valve development^32,41^ as a proxy for investigating the impact of variants associated with dysplastic left or right cushions as these share a common mechanism^17^. More generally, it is possible that leaflet dysplasia or hyperplasia could be modelled if all leaflets share a common maturation processes.

There are numerous benefits to studying valvulogenesis in zebrafish: temporal reproducibility of the process, accessibility of the key stages, external development for live imaging and that this is all possible prior to reaching the stage at which studies become regulated (typically 5dpf). Whilst previously, the majority of developmental studies on valve developmental has occurred in mice and then shown to be relevant in humans and other species such as zebrafish, it is now apparent that more developmental work can be done principally in zebrafish and confirmed in mice, greatly reducing the use of animals in fundamental research.

In summary, we have shown that the two leaflets of the zebrafish arterial valve are established by the same developmental mechanism as the mouse and human intercalated leaflets, distinct to that of atrioventricular valve formation. This opens up new avenues for using zebrafish in understanding human congenital heart disease through functional testing of human gene variants.

## Acknowledgements

We thank Rashmi Priya, Elizabeth Patton, Emily Noël and Tanya Whitfield for provision for zebrafish lines. We are grateful to the Newcastle University Aquarium Technical team for their support and assistance in this work. We apologise to authors whose work could not be included due to limitations on references.

