## Supplemental files for "Zebrafish arterial valve development occurs through direct differentiation of second heart field progenitors"

*Supplemental Figures 1-6*

*Movies 1-4*

*Supplemental Methods*

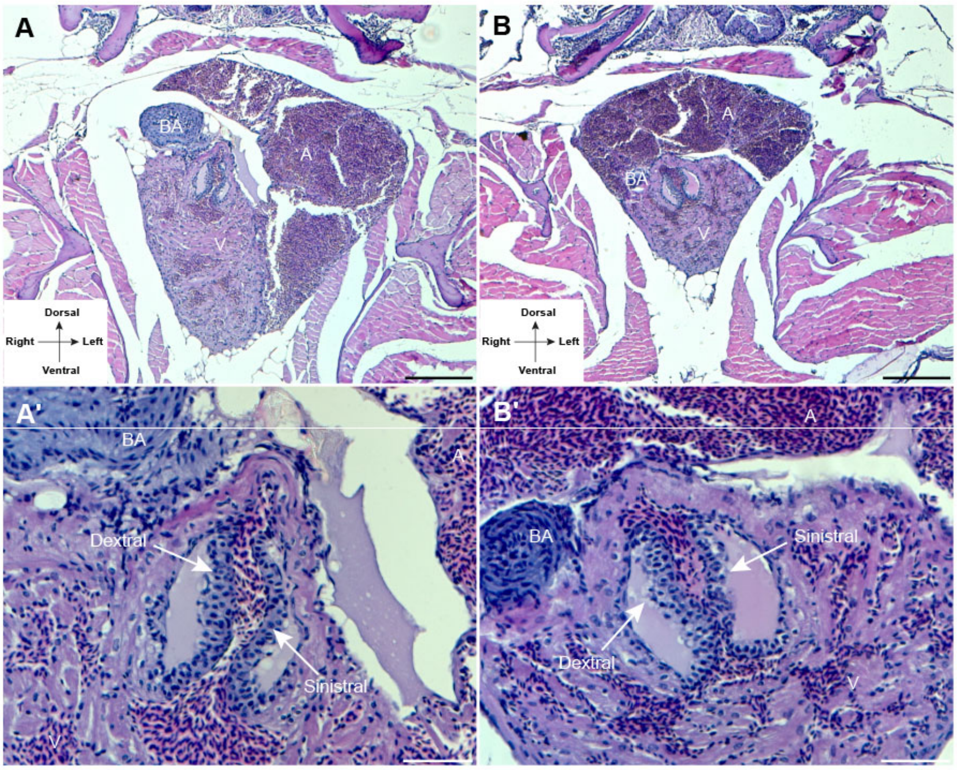


**Supplemental Figure 1. Zebrafish arterial valve leaflets are positioned laterally. (A-A’)** Anterior view of 117dpf (Standard Length: 2.28cm) female zebrafish stained with Haematoxylin and Eosin showing the position of heart within the body cavity (A) and the orientation of the leaflets (A’). **(B-B’)** Anterior view of 117dpf (Standard Length: 1.88cm) male zebrafish stained with Haematoxylin and Eosin showing the position of heart within the body cavity (B) and the orientation of the leaflets (B’). The line of apposition of the arterial valve leaflets is along the antero-posterior axis, and the leaflets are predominantly located on the left (sinistral) and right (dextral). V: Ventricle, A: Atrium, BA: Bulbous Arteriosus. Dorsal: up. Scale bars: A, B: 200µm, A', B’: 50µm.

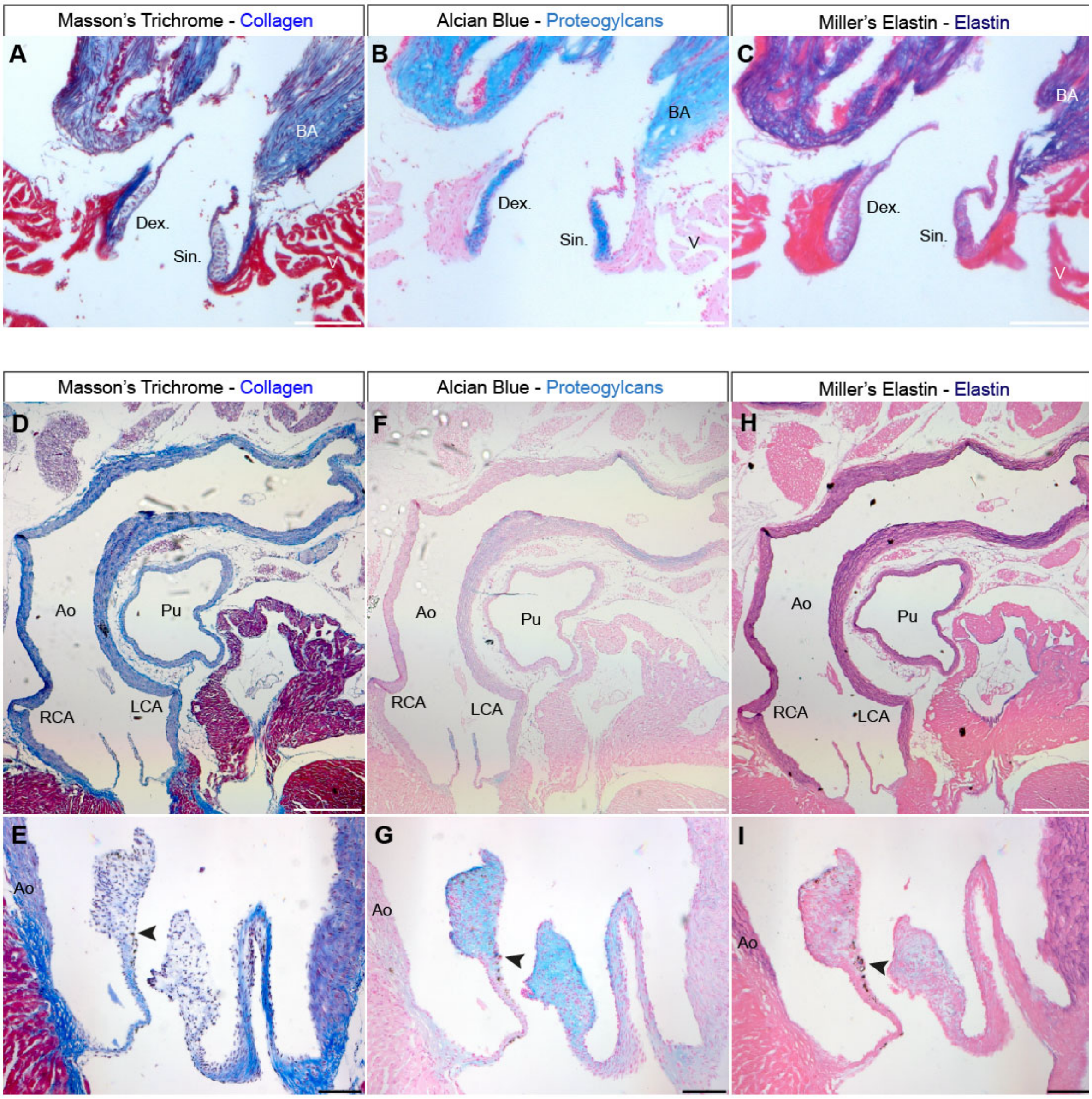


**Supplemental Figure 2. Localisation of ECM components in the adult zebrafish and mouse arterial pole.**

**(A)** Masson's Trichrome stained longitudinal section of the adult zebrafish arterial pole (wide field view of Fig. 1G). Collagen is present in the Bulbous arterious and continuous with the arterial root. **(B)** Alcian Blue stained longitudinal section of the adult zebrafish arterial pole (wide field view of Fig. 1I). The Bulbous arteriosus is rich in sulphated proteogylcans, with some staining present in the distal-most part of the arterial root. **(C)** Miller's Elastin stained longitudinal section of the adult zebrafish arterial pole (wide field view of Fig. 1K). Elastin is abundant in the Bulbous arteriosus and continuous with the arterial root. **(D)** Masson's Trichrome stained longitudinal section of the arterial pole of a P90 mouse. Collagen is present in the great vessels and the root of the aortic valve. Wide field view of Fig. 1L. **(E)** Oblique cut of left and right aortic valve leaflets of P90 mouse stained with Masson's Trichrome. Collagen is present in the root, hinge and base of the leaflets and more strongly than in the wall of the aorta. **(F)** Alcian Blue stained longitudinal section of the arterial pole of a P90 mouse. Sulfated proteoglycans are present at low levels in the great vessels. Wide field view of Fig. 1N. **(G)** Oblique cut of left and right aortic valve leaflets of P90 mouse stained with Alcian Blue. Sulfated proteoglycans are found concentrated in the tips of the leaflets in a reciprocal pattern to Collagen and present at low levels in the hinge. Very little stain is present in the wall of the aorta. **(H)** Miller's Elastin stained longitudinal section of the arterial pole of a P90 mouse. Mature elastin fibres are clearly visible in the wall of the aorta and pulmonary artery. Wide field view of Fig. 1P. **(I)** Oblique cut of left and right aortic valve leaflets of P90 mouse stained with Miller's Elastin. Mature elastic fibres are present in the wall of the aorta, with diffuse staining in the tips likely reflecting immature elastin. Elastin is absent from the hinge and base of the leaflets. Neural-crest derived melanocytes are observable in some sections (arrowheads, E, G, I). BA: Bulbous arteriosus, V: Ventricle, Dex: Dextral leaflet, Sin: Sinistral leaflet, Ao: Aorta, Pu: Pulmonary artery, RCA: Right coronary artery, LCA: Left coronary artery. Anterior: up. Scale bars: A-C: 100μm. D, F, H: 500μm. E, G, I: 50μm.

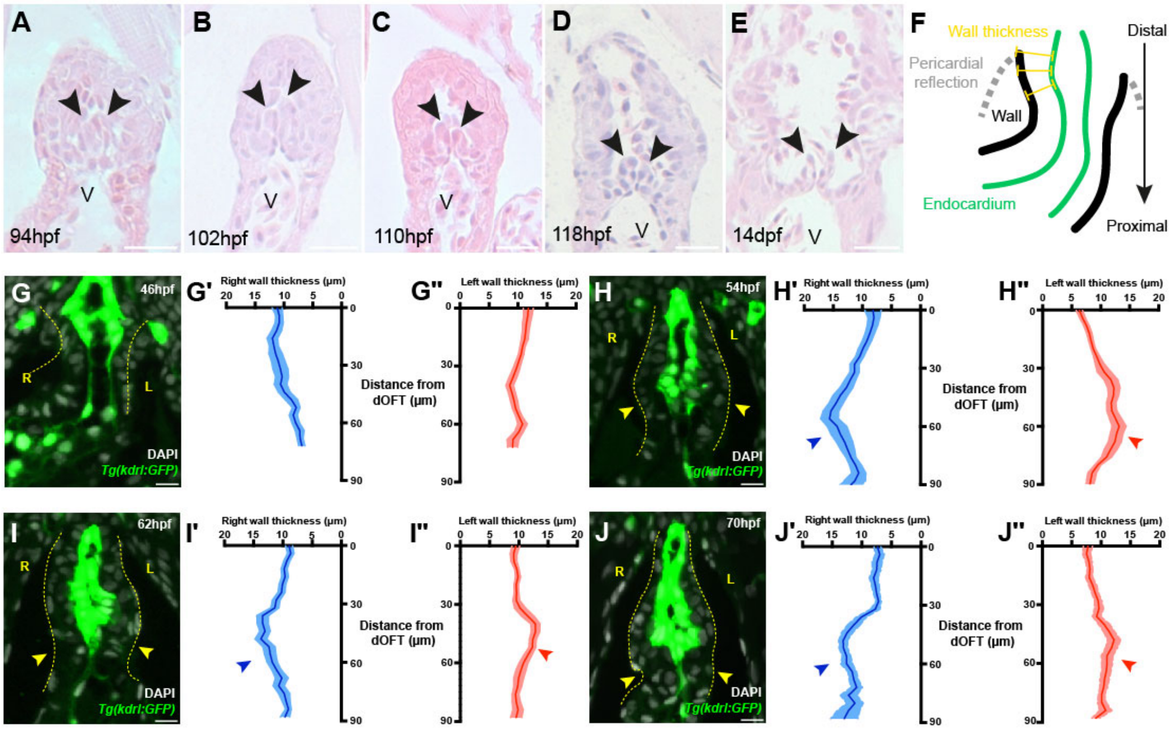


**Supplemental Figure 3. Arterial valve primordia formation occurs between 2-3dpf.**

**(A-E)** Representative Haematoxylin and Eosin-stained midline section along the long axis of the OFT at 94hpf (A) (n=5), 102hpf (B) (n=6), 110hpf (C) (n=5), 118hpf (D) (n=5) and 14dpf (E) (n=5). Two leaflets are visible (arrowheads), which have begun to remodel between 5-14dpf and appear thinner. **(F)** Schematic of approach to measure wall thickness. The perpendicular distance (yellow) between the inner edge of the endocardium (marked by Tg(kdrl:GFP), green) and the outer edge of the wall (black) is measured with the pericardial reflection (grey) defined as the distal most point of the OFT. **(G-G")** Quantification of wall thickness at 46hpf. No primordia are present at this stage. Yellow line denotes edge of wall (n=11). **(H-H")** Quantification of wall thickness at 54hpf. Increased wall thickness, representing arterial primordia are visible. Yellow line denotes edge of wall, yellow arrowheads in (H) denote primordia in (H'-H") (n=8). **(I-I")** Quantification of wall thickness at 62hpf. Increased wall thickness, representing arterial primordia are visible. Yellow line denotes edge of wall, yellow arrowheads in (I) denote primordia in (I'-I") (n=11). **(J-J")** Quantification of wall thickness at 70hpf. Increased wall thickness, representing arterial primordia are visible. Yellow line denotes edge of wall, yellow arrowheads in (J) denote primordia in (J'-J") (n=10). V: Ventricle, dOFT: distal outflow tract, R:

morphological right, L: morphological left. Anterior: up. G-J: thickness is mean ± S.E.M. Scale bars: A-E: 20μm, G-J: 10μm.

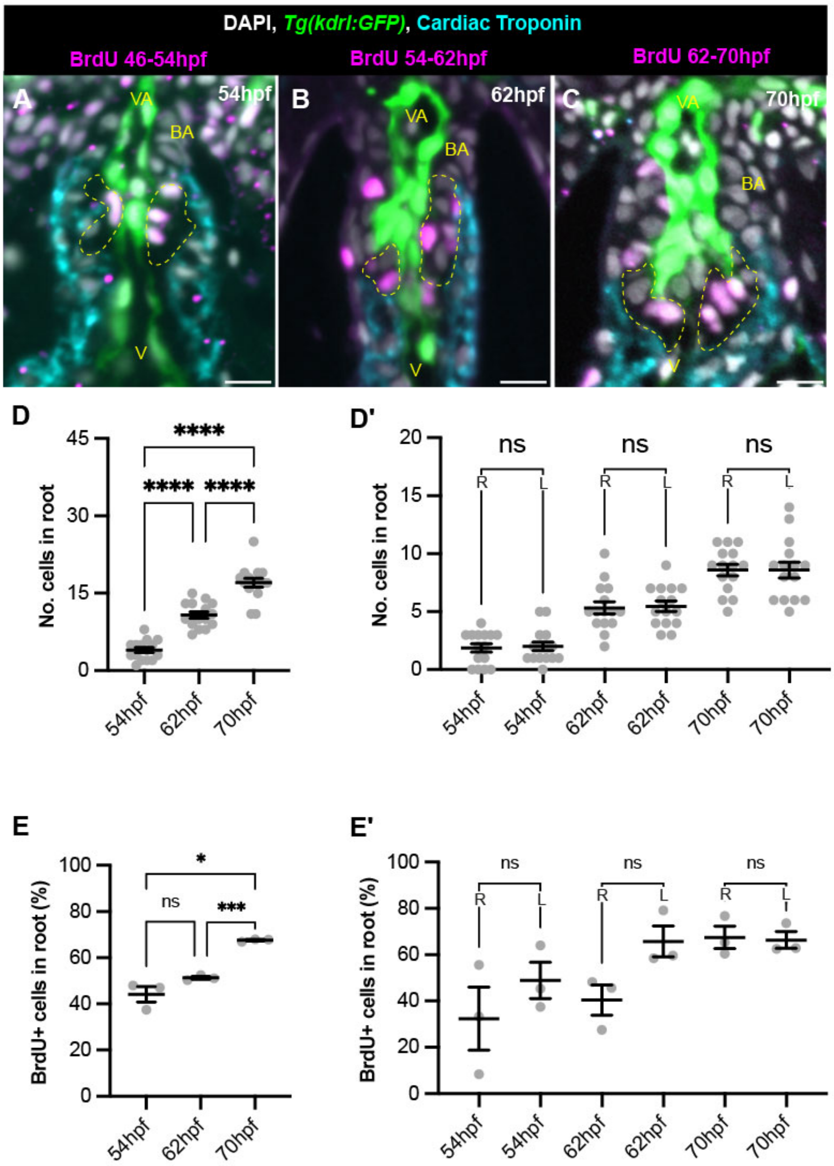


**Supplemental 4. Proliferation in the arterial valve primordia accompanies increase in cell number**

**(A)** Representative midline resin section of *Tg(kdrl:GFP)* (green) embryo incubated in BrdU between 46-54hpf, fixed at 54hpf and stained for BrdU (magenta) and Cardiac Troponin (cyan). Region of quantification (primordia) denoted by yellow lines. **(B)** Representative midline resin section of *Tg(kdrl:GFP)* embryo incubated in BrdU between 54-62hpf, fixed at 62hpf and stained for BrdU and Cardiac Troponin. Region of quantification (primordia) denoted by yellow lines. **(C)** Representative midline resin section of *Tg(kdrl:GFP)* embryo incubated in BrdU between 62-70hpf, fixed at 70hpf and stained for BrdU and Cardiac Troponin. Region of quantification (primordia) denoted by yellow lines. **(D)** Quantification of total cell number in the arterial valve primordia between 54-70hpf, cell number significantly increases over the course of development. **(D')** Breakdown of quantification in (D) into right (R) and left (L) there are no differences between the left and right primordia (N=3, 5 fish per N, n=15). (E) Quantification of percentage of BrdU positive cells in the arterial valve primordia between following BrdU pulses, levels of proliferation remain relatively constant over the course of development. **(E')** Breakdown of quantification in (E) into right (R) and left (L) there are no differences between the left and right primordia (N=3, 5 fish per N). V: Ventricle, VA: Ventral Aorta, BA: Bulbous Arteriosus. Scale bars: 10µm. D-E': Mean ± S.E.M, Brown-Forsythe and Welch ANOVA with multiple comparisons. *: $p<0.05$, ***: $p<0.001$, ****: $p<0.0001$, ns: not significant.

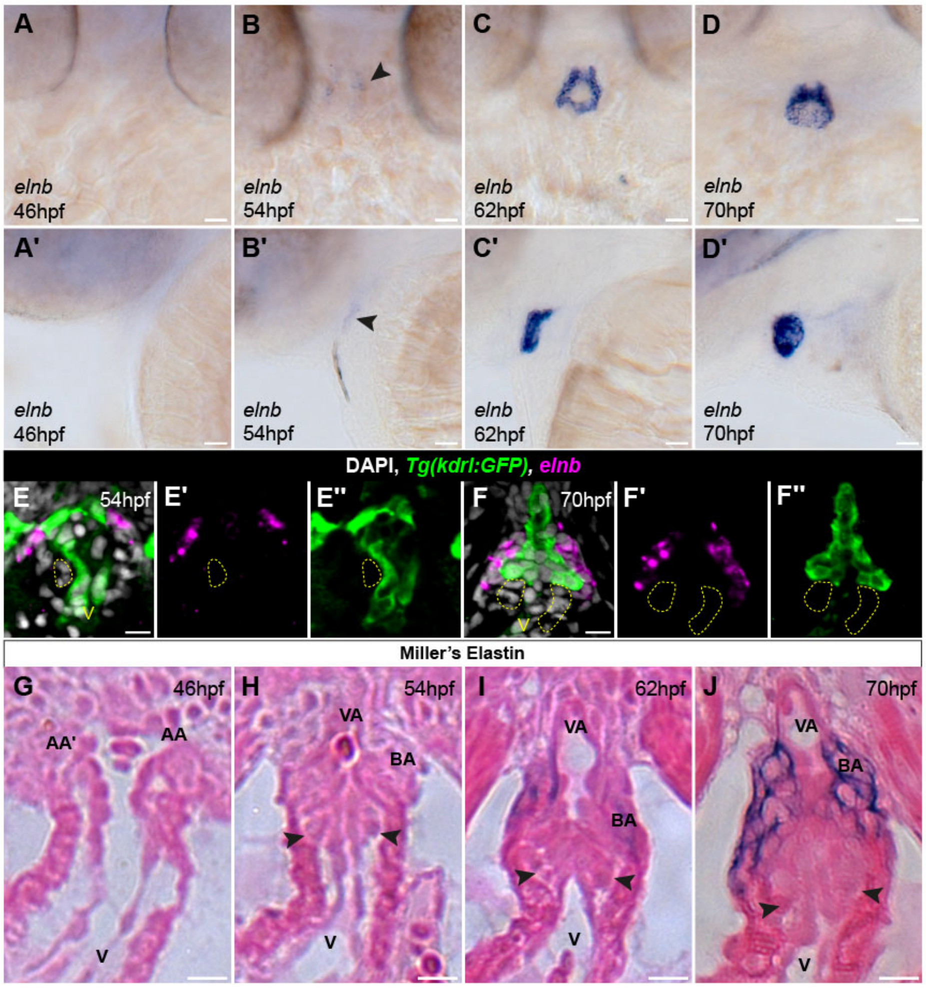


**Supplemental Figure 5. *elnb* expression coincides with primordia appearance**

**(A-D')** mRNA *in situ* hybridisation analysis of *elnb* expression during arterial valve primordia formation. *elnb* is not expressed at 46hpf (A-A') (n=17), initiates at 54hpf (B-B' arrowheads) (n=17) and the domain expands between 62hpf-70hpf (C-D') (62hpf n=17, 70hpf n=17). **(E-E")** Midline resin section of fluorescent mRNA *in situ* hybridisation of *elnb* (magenta, E') in *Tg(kdrl:GFP)* (green, E") embryo at 54hpf counterstained with DAPI (white). *elnb* is not expressed in the cells of the primordia (yellow) but is expressed at the distal-most aspect of the OFT (n=7). **(F-F")** Midline section of fluorescent mRNA *in situ* hybridisation in of *elnb* (magenta, F') in *Tg(kdrl:GFP)* (green, F") embryo at 70hpf counterstained with DAPI (white). *elnb* is not expressed in the cells of the primordia (yellow) but is expressed at the distal-most aspect of the OFT (n=7). **(G-J)** Midline resin sections of embryos stained for elastin fibres by Miller's Elastin between 46-70hpf. No elastin fibres are present at 46-54hpf (G, H)

(46hpf, n=13. 54hpf, n=12). Fibres are present at 62hpf (I) (n=11) and by 70hpf (J, n=12), the majority of the BA has elastin fibres, but these are not present in the primordia (arrowheads in H-J). A-D, E-J : Ventral views, A'-D': Lateral views, anterior left. AA: Aortic arches, V: Ventricle, VA: Ventral Aorta, BA: Bulbous Arteriosus. Scale bars: 10μm.

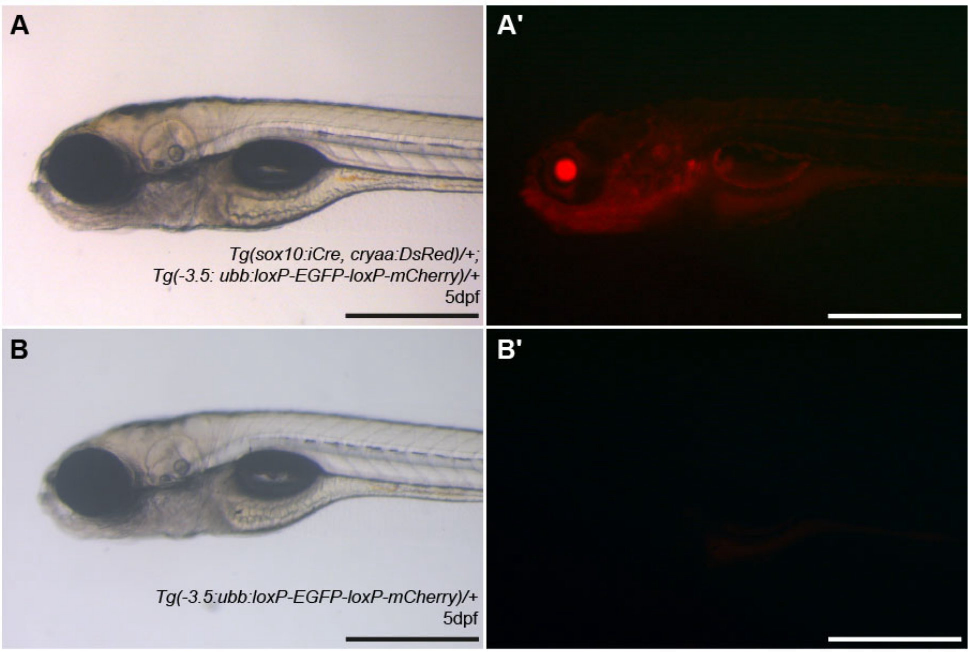


**Supplemental Figure 6. Characterisation of *Tg(sox10:iCre, cryaa:DsRed2)*$^{n6}$ line**

**(A)** Brightfield image of live *Tg(sox10:iCre, cryaa:DsRed2)*/+; *Tg(-3.5:ubb:loxP-EGFP-loxP-mCherry)/*+ at 5dpf. **(A')** Same fish from (A) viewed under fluorescence with DsRed filter. Recombination is present in neural crest-derived structures (craniofacial). Signal in the eye is the transgenesis marker. **(B)** Brightfield image of live *Tg(-3.5:ubb:loxP-EGFP-loxP-mCherry)*/+ at 5dpf. **(B')** Same fish from (B) viewed under fluorescence with DsRed filter. No recombination is visible. Anterior: left. Scale bar: 50µm

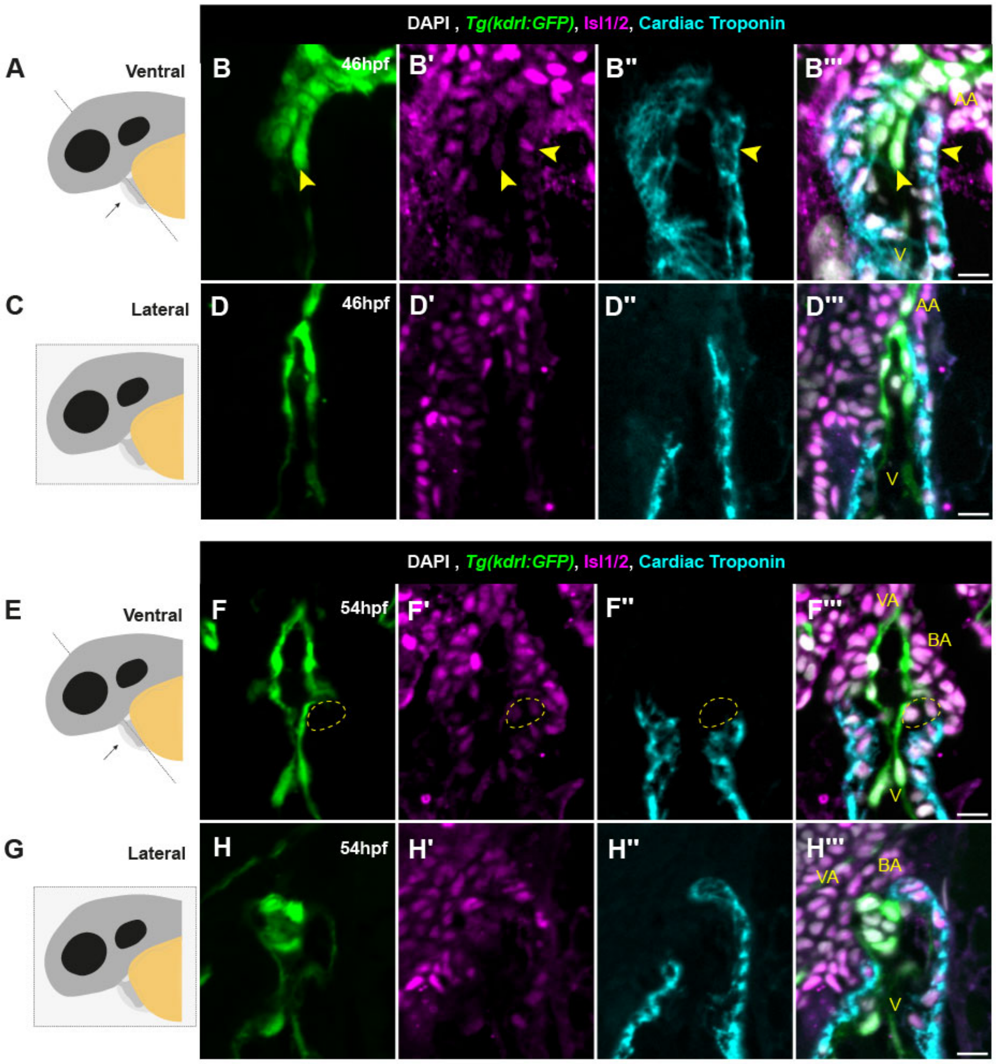


**Supplemental Figure 7. Characterisation of the transition zone in zebrafish. (A)** Schematic showing plane of section for ventral view of the heart. **(B-B''')** Ventral view of arterial pole of the heart at 46hpf, with the endocardium marked by Tg(kdrl:GFP) (B, green), Isl1/2+ cells in magenta (B') and myocardium marked by Cardiac Troponin (B'' cyan). Both cardiomyocytes and endocardium at the distal end of the OFT still maintain Isl1/2 expression (arrowheads), confirming the presence of the arterial pole transition zone. (B''') (n=14). **(C)** Schematic showing plane of section for lateral view of the heart. **(D-D''')** Lateral view of arterial pole of the heart at 46hpf, with the endocardium marked by Tg(kdrl:GFP) (D, green), Isl1/2+ cells in magenta (D') and myocardium marked by Cardiac Troponin (D'', cyan). The myocardium extends more anteriorly on the ventral surface of the heart. Both cardiomyocytes and endocardium at the distal end of the OFT still maintain Isl1/2 expression (D''') (n=9). **(E)** Schematic showing plane of section for ventral view of the heart. **(F-F''')** Ventral view of arterial pole of the heart at 54hpf, with the endocardium marked by Tg(kdrl:GFP) (F, green), Isl1/2+ cells in magenta (F') and myocardium marked by Cardiac Troponin (F'' cyan). Both cardiomyocytes and endocardium at the distal end of the OFT still maintain Isl1/2 expression. Cells of the forming primordia (yellow region) and the arterial domain distal to the myocardium are positive for Isl1/2 (F''') (n=14). **(G)** Schematic showing plane of section for lateral view of the heart. **(H-H''')** Lateral view of arterial pole of the heart at 54hpf, with the endocardium marked by Tg(kdrl:GFP) (H, green), Isl1/2+ cells in magenta (H') and myocardium marked by Cardiac Troponin (H'', cyan). The myocardium extends more anteriorly on the ventral surface of the heart. Both cardiomyocytes and endocardium at the distal end of the OFT still maintain Isl1/2 expression, as well as the arterial domain distal to the myocardium (H''') (n=10). AA: Aortic arch, BA: Bulbous Arteriosus, V: Ventricle, VA: Ventral Aorta. Scale bars: 10μm.

**Movie 1. Arterial valve primordia at 62hpf**

Representative movie of Tg(kdrl:GFP) at 62hpf (Fig 3I-I’) at half live speed. Two primordia are visible between the Ventricle and Bulbous Arteriosus. BA: Bulbous Arteriosus, V: Ventricle, A: Atrium.

**Movie 2. Arterial valve primordia at 70hpf**

Representative movie of Tg(kdrl:GFP) at 70hpf (Fig 3J-J’) at half live speed. Two primordia are visible between the Ventricle and Bulbous Arteriosus. BA: Bulbous Arteriosus, V: Ventricle, A: Atrium.

**Movie 3. Arterial valve primordia at 78hpf**

Representative movie of Tg(kdrl:GFP) at 78hpf (Fig 3K-K’) at half live speed. Sinuses have begun to form in the two primordia are visible between the Ventricle and Bulbous Arteriosus. BA: Bulbous Arteriosus, V: Ventricle, A: Atrium.

**Movie 4. Arterial valve primordia at 86hpf**

Representative movie of Tg(kdrl:GFP) at 86hpf (Fig 3K-K’) at half live speed. Sinuses have formed in the two primordia, defining the leaflets of the arterial valve between the Ventricle and Bulbous Arteriosus. BA: Bulbous Arteriosus, V: Ventricle, A: Atrium.

### Supplemental Methods

#### Zebrafish lines and general husbandry

The following previously described lines were used: *Tg(kdrl:GFP)*$^{la116}$ (1), *Tg(kdrl:Cre)*$^{s898}$ (2), *Tg(-3.5:ubb:loxP-EGFP-loxP-mCherry)*$^{cz1701}$ (3), *Tg(fli1a:AC-TagRFP)*$^{sh511}$ (4), Tg(myl7:EGFP)$^{twu34}$ (5), *Tg(kdrl:HsHRAS-mCherry)*$^{s916}$ (6), *tbx1*$^{tm208}$ (7–9), *vangl2*$^{m209}$ (9,10). *Tg(kdrl:GFP)*$^{la116}$ was maintained in a Capser (*roy*$^{a9/a9}$*; mitf*$^{aw2/aw2}$) background (11), all other lines were maintained in a mixed AB background. *tbx1*$^{tm208}$ genotyping was carried out as previously described (8). *vangl2*$^{m209}$ genotyping was carried out as previously described (12). *Tg(kdrl:Cre)*$^{s898}$ was maintained by breeding Cre+ males with WT(AB) females (13,14).

Embryos were obtained from natural pairwise mating of adults, maintained at 28.5 degrees C in E3 medium (5 mM NaCl, 0.17 mM KCl, 0.33 mM CaCl2, 0.33 mM MgSO4) and staged according to Kimmel 1995 (15). Where necessary, between 22-24 hpf, embryos were transferred into E3 medium containing 0.003% 1-phenyl-2-thiourea (PTU, Sigma P7629) to prevent pigment formation, aiding imaging. Embryos were fixed in 4% Paraformaldehyde (PFA, P6148 Sigma) in 1x Phosphate Buffered Saline (PBS, Oxoid BR0014G) overnight at 4 degrees C. The next day embryos were washed in PBST (1xPBS with 0.2% Tween (Sigma P2287)) and serially dehydrated to 100% MeOH and stored at -20C or washed in PBS and serially dehydrated to 70% EtOH and stored at room temperature. For mutant analyses, embryos were rinsed from fixative into PBST, the caudal fin taken for gDNA extraction and genotyping as above. Once genotyped, WT siblings and mutant embryos were collated and dehydrated separately.

**Collection of adult zebrafish tissue**

Adult zebrafish hearts were manually dissected following terminal anaesthesia using Tricaine methanesulfonate (MS222, Merck E10521) and destruction of the brain. Hearts were briefly incubated in room temperature Ringer's solution with 10mM Calcium Chloride (116mM NaCl, 2.9mM KCl, 10mM $CaCl_2$, 5mM HEPES), before overnight fixation in 4% PFA at 4 degrees C. Hearts were washed three times in PBS at room temperature, and then serially dehydrated to 70% EtOH for long term storage at room temperature.

Adult zebrafish were culled by terminal anaesthesia (MS222, Merck E10521) and destruction of the brain, with images taken prior to fixation for measurement of standard length (16). Whole fish were fixed at in 4% PFA at 4C for 4 nights, washed 3 times in PBS at room temperature, decalcified in 0.5M EDTA pH8.0 (Merck, 27285) with agitation at 4C for 4 nights, and then serially dehydrated to 70% EtOH for long term storage at room temperature.

**Establishment of *Tg(sox10:iCre, cryaa:DsRed2)* line**

The previously described *sox10:iCre,cryaa:DsRed2* plasmid (17) was injected together with *tol2* mRNA (18) and Phenol Red (Sigma, P0290) into the yolk of 1-cell stage embryos obtained from in-crossing WT(AB) adults. Each embryo was injected with 33.3pg of plasmid, 33.3pg of tol2 mRNA and 25% Phenol Red. Healthy F0 embryos were screened at 4dpf for the presence of DsRed2+ eyes (marker of transgenesis) to enrich for plasmid integration and raised to adulthood. Adult F0 were outcrossed to WT(AB), embryos with DsRed+ eyes at 4dpf were raised to adulthood (F1). A single F1 adult male was selected to establish the line at F2, based on 50% transmission of DsRed+ eyes at 4dpf (representing a single insertion). All experiments were carried out using at least F3. The is designated *Tg(sox10:iCre, cryaa:DsRed2)*$^{n6}$ and is maintained by breeding Cre+ males with WT(AB) females (13,14).

**Lineage tracing**

Adult male heterozygous *Tg(kdrl:Cre)*$^{s898}$ or *Tg(sox10:iCre, cryaa:DsRed2)*$^{n6}$ were outcrossed to *Tg(-3.5:ubb:loxP-EGFP-loxP-mCherry)*$^{cz1701}$ females to ensure tissue specificity of Cre activity (13,14). Embryos positive for the reporter were fixed at 70hpf, washed three times in PBST and genotyped for Cre. *Tg(kdrl:Cre)*$^{s898}$ using primers Forward: 5'-AACGAGTGATGAGGTTCGCAAGAACC-3' and Reverse: 5'-AAATCAGTGCGTTCGAACGCTAGAG-3' with GoTaq G2 (Promega): 35 cycles, 30s extension time, 55C annealing temperature. *Tg(sox10:iCre, cryaa:DsRed2)*$^{n6}$ using primers Forward: 5'- GCAGCCACACCATTCTTTCT-3' and Reverse: 5'-GAAGACCATCCAACAGCACC-3' with GoTaq G2: 35 cycles, 30s extension time, 58C annealing temperature. Embryos were split into Cre+ve and Cre-ve and serial dehydrated to 100% MeOH at -20C. Cre-ve embryos showed no recombination (data not shown).

**mRNA *in situ* hybridisation**

For chromogenic mRNA *in situ* hybridisation, embryos were fixed overnight in 4% paraformaldehyde (PFA, Cell Signalling Technology 12606) and steps were carried out as previously described (19). Fluorescent mRNA in situ was performed using the APExBIO Tyr-Cy3 TSA Kit (Stratech, K1051-APE) and carried out as previously described (20), the *Tg(kdrl:GFP)* signal was boosted post-ISH by immunohistochemistry as described above, but with only 30 mins of blocking. FISH samples were rapidly dehydrated before resin embedding.

**Resin embedding of zebrafish embryos**

For resin sectioning, embryos and adult hearts were dehydrated in 100% EtOH (this is performed rapidly for embryos with fluorescent staining, see (21)). Tissue was moved from EtOH into Hardener 1 (Technovit 8100), allowed to settle and then rinsed twice in Hardener 1, and incubated at 4C for at least 2hrs. For embedding, Hardener 2 was mixed with fresh Hardener 1 according to manufacturers instructions, tissue was transferred into H1-H2 mix, oriented and sealed in an airtight container at 4C overnight. Where necessary, blocks were cut and re-embedded for specific orientations.

Adult hearts were cut at 7µm sections and embryos cut at 4µm sections using a RMC MT990 Microtome. H&E of resin sections was carried out as previously described (21) and allowed to dry. H&E and embryonic Miller's were mounted in Histomount. Resin sections of immunohistochemistry where mounted in 1:1 mix PBS-Vectashield with DAPI (Vector labs H-1200), or where Cardiac Troponin was revealed using Alexa350, 1:1 PBS-Vectashield (Vector labs H-1000) with VectaShield with DAPI diluted to 0.3%. Details of quantifications are given in the supplemental methods.

**Quantification of wall thickness**

Midline sections of embryos, where a lumen is visible extending from the ventricle to the connection of the heart with the surrounding mesoderm were selected. In Fiji (22,23), using an adapted add_mark_every_x_microns.ijm macro (https://gist.github.com/mutterer/95cc6f027080674af68ca65fae8d1051) the length of the left and right *Tg(kdrl:GFP)* signal at the distal most point of the heart (defined as 0 for measurements) was marked and subdivided into 4µm sections. Using the DAPI channel, the outer surface of the wall of the heart was traced using the freehand line tool and the perpendicular distance from the endocardium to the outer edge of the wall was measured. The myo-arterial boundary was defined as the first 4µm subdivision of the *Tg(kdrl:GFP)* signal in which the wall was positive for Cardiac Troponin signal. For embryos where more than one section was measured, an average of the wall thickness across the sections was taken.

**Quantification of primordia cell number, proliferation, OFT length and *elnb* domain**

In Fiji, sections through the OFT were selected on the basis of a clear lumen with continuous endocardial transgene signal from the distal-most point of the OFT to the ventricle. Using the myocardial signal, a straight line was drawn across the distal-most signal – defining the arterial primordia as cells below this line that were present between the myocardium and endocardium but did not express either of these markers. Cells were counted in alternate sections.

For BrdU analysis, cells were defined as BrdU+ or BrdU- within the arterial valve primordia (as defined above). For each individual embryo (n), average BrdU+ cells in the primordia were quantified (as above). A single average was calculated from the 5 embryos of one treatment group (N), this was repeated twice, giving 3 experimental replicates, each consisting of 5 embryos.

In the same sections, arterial domain length was measured using the freehand line tool, starting at the distal-most point of the OFT and ending at the myocardial signal, an average of the arterial OFT length across all midline sections was taken.

The shape of the *elnb* signal was traced using the freehand selection tool and shape parameters measured. Embryos were genotyped post image acquisition.
